# Genetic admixture between East and West European Gravettian-associated populations in Western Europe before the Last Glacial Maximum

**DOI:** 10.1101/2025.09.26.678744

**Authors:** Pere Gelabert, Susanna Sawyer, Olivia Cheronet, Vanessa Villalba-Mouco, Victoria Oberreiter, Manuel Ramón González-Morales, Lawrence G. Straus, Igor Gutiérrez-Zugasti, David Cuenca-Solana, Diego Gárate, Ana B. Marín-Arroyo, Henry de Lumley, José-Miguel Tejero, Christian Normand, Joëlle Darricau, Michaela Lucci, Alessia Nava, Francesco Genchi, Donato Coppola, Chiara La Marca, Thomas Higham, Maddalena Giannì, Laura G. van der Sluis, Carla Gómez-Montes, Michelle Hämmerle, Brina Zagorc, Florian Exler, Florian Brück, Stefan Franz, Fernanda Tenorio Cano, Kristin Stewardson, Lijun Qiu, Mareike Stahlschmidt, Alfredo Coppa, David Reich, Ron Pinhasi

## Abstract

Modern humans first settled in Europe at least 45,000 years ago. However, limited genomic data from individuals dating between 45,000 and 20,000 years ago still restricts our understanding of population dynamics and admixture during the Upper Palaeolithic. Before the Last Glacial Maximum (LGM, 26.5–19 cal kya), Gravettian culture-associated populations were widespread and genetically diverse, comprising at least two distinct genetic groups, referred to as the Fournol and Věstonice clusters. We present genome-wide data from three Gravettian-associated individuals: two from cave sites in the Franco-Cantabrian region (Chufín and Isturitz) and one from Italy (Ostuni1b). These data reveal previously undetected gene flow linking the ancestry of 34,000-year-old individuals from Sungir (Russia) to Gravettian individuals from Western Europe, challenging the prevailing model of population continuity from the Aurignacian to the Solutrean. As osseous remains are scarce for this time period, DNA from sediments deposited by ancient humans opens a new possibility to obtain genomic data. We thus examine sedimentary DNA from Solutrean Layer 122 at El Mirón Cave (Cantabria, ∼22,000 cal BP), recovering approximately 16,000 human SNPs, among the highest yields reported from a Palaeolithic context. Generating these data required over 1.15 billion sequencing reads, illustrating both the potential of sediment DNA for autosomal analysis and the technical challenges of the approach.

## Introduction

The Gravettian (ca. 34–25 ka cal BP)^1,2^ was a distinctive, widespread archaeological culture that existed before and at the outset of the Last Glacial Maximum in Europe. Despite widespread similarities in lithic and osseous technologies, burial practices, and portable artworks (especially the so-called “Venus” figurines), the Gravettian was not an entirely homogeneous culture. Through the study of material remains, up to nine regional groups have been proposed to have existed in this period^3^. The available Gravettian pre-LGM genomes from central and western Europe are scarce (Vêstonice 13,15,16,43 (Czechia), Pavlov1 (Czechia), Paglicci12 (Italy), six Goyet individuals (Belgium), Ostuni1 (Italy), Ostuni2 (Italy), Mollet III (Spain), Reclau-Viver (Spain), La Rochette (France), Fournol (France), two Krems-Wachtberg (Austria) individuals and Ormesson (France)) ^4–6^, which limits our understanding of the extent of genetic isolation among these defined clusters, temporal dynamics during the Gravettian, and the sequence and timing of ancestry changes leading to the Solutrean (Western Europe, 24.5–20 ka cal BP) and Epigravettian (Italy and Southeastern Europe, 24–12 ka cal BP) cultures and peoples. The genetic data available to date show the existence of at least two different genetic ancestry clusters in Europe: a western ancestry, referred to as the Fournol cluster, represented by the shared affinity to the Aurignacian-associated genome of GoyetQ116-1 (Belgium) and the Gravettian-associated genome of Fournol (Southern France)^5,6^, and a second cluster, referred to as the Vêstonice cluster, as indicated by the genomes recovered from Dolní Vêstonice’s Gravettian levels (Czechia) and the rest of Central European and Italian Gravettian-associated individual genomes^5,6^. Instances of admixture between these two distinct Gravettian ancestries are rare. However, one notable case is a group from Goyet in Belgium (27 ka cal BP, Late Gravettian), which shows a mixed ancestry between the Věstonice and Fournol genetic clusters. This admixture suggests an east-to-west expansion of Věstonice-associated ancestry occurred between the Early/Middle and Late Gravettian, resulting in a longitudinal cline of gene mixing between these previously distinct pre-LGM populations ^6^. To date, none of these signals of admixture with other eastern European sources has been identified in South or Western Europe before the LGM.

Recent advances in DNA retrieved from ancient sediment layers (sedaDNA) have demonstrated the capacity to retrieve mammalian mitochondrial data from sediment samples ^7–9^. Some studies have also recovered nuclear DNA, which enables the study of population histories, albeit with limited resolution^8,10^. Harnessing sedaDNA from successive deposits can enable fine-scaled diachronic analysis (but see ^11,12^), which cannot be achieved from the paleogenomic study of Palaeolithic human remains, as these are scarce ^7,13^. Even with the screening of unidentified bone fragments via collagen fingerprinting^14^, such approaches rarely provide a fine-grained perspective ^7,13^. This approach is especially relevant for archaeologically defined cultural entities like the Gravettian and Solutrean, that persisted for several thousands of years, but for which genome-wide data are available for only 29 individuals ^4–6^. Our recent study of El Mirón’s Solutrean sediments using an in-solution mitochondrial DNA (mtDNA) capture approach has produced the most substantial mitochondrial results for humans and a range of mammalian species in Paleolithic context^7^. Building on these results, we developed a strategy to recover a sufficient number of informative nuclear single-nucleotide polymorphisms (SNPs) from ElMiron14 sediment sample (Level 122, Solutrean), which enabled its analysis through standard paleogenomic ancestry analyses, including Principal Components Analysis (PCA) and *f-statistics*. In addition, we generated genome-wide data from skeletal remains of three further Gravettian individuals: a human tooth from Chufín Cave in Cantabria, a long bone from Isturitz Cave in the French Basque Country, and a fetal tibia from Ostuni Cave (Ostuni1b), Southern Italy (27,810–27,430 cal BP)^5^. Combined with previously published data, this dataset provides a meaningfully richer genomic record of Pleistocene humans for the Gravettian and Solutrean periods. Our study also highlights key challenges in sedaDNA studies aimed at investigating genomic change, while revealing previously undetected regional genetic patterns that transcend strict chronological boundaries.

## Results

### The generation of low-coverage sedaDNA genome-wide data

We aimed to obtain genome-wide data from the ElMiron14 sediment sample (Supplementary Figures 1 and 2), one of the best preserved samples from our previous mtDNA analysis of sediments from this site^7^. To generate sufficient data for genome-wide analysis, we produced five libraries from two extracts of ElMiron14^7^, indexing and sequencing each of the libraries an average of ten times (in some cases following in-solution enrichment), thereby producing 1,028,866,146 sequences in total (Supplementary Data 1). Of the 101 individual sequencing attempts, two-thirds (68) were performed after in-solution enrichment with the TWIST 1.4M^15^ human nuclear capture kit, and one-third (33) were performed without enrichment (shotgun sequencing) (Supplementary Information). After filtering, an average of 3,018 reads per library were assigned to humans in the capture experiments, compared to 2,779 in the shotgun libraries. The results indicate no significant difference in the number of human reads recovered between the two sequencing strategies after PCR duplicates were removed and reads were filtered with MEGAN. *(W=1040.5, P=0.636)* (Figure 1B, Supplementary Information). The comparison, however, reveals a difference in the number of SNPs recovered on the 1.4M^16^ capture versus those recovered on the same 1.4M range using the shotgun approach. The combined 1.4M capture libraries produced 48,384 informative positions, compared to just 1,515 from shotgun sequencing, a 32-fold increase in data recovery (but 128-fold if sequencing yields are taken into consideration (Supplementary Data 1). Theoretical expectations, considering that the human genome comprises approximately 3 billion base pairs (bp), the 1.4 million (1.4M) SNP capture reagent targets roughly one in 2,143 bp of the genome. Assuming an average fragment length of 60 bp (Supplementary Information), this enrichment strategy theoretically increases the probability of sequencing a targeted site by approximately 36-fold compared to standard shotgun sequencing. The capture approach exceeded those expectations. We obtained 11,210 mtDNA unique reads from these libraries by capturing the mtDNA genome (Supplementary Information, Supplementary Data 2). All the libraries show deamination and diversity classified within the mtDNA haplogroup U, coincident with the previous classification of the ElMiron14 sample ^7^.

**Figure 1.**
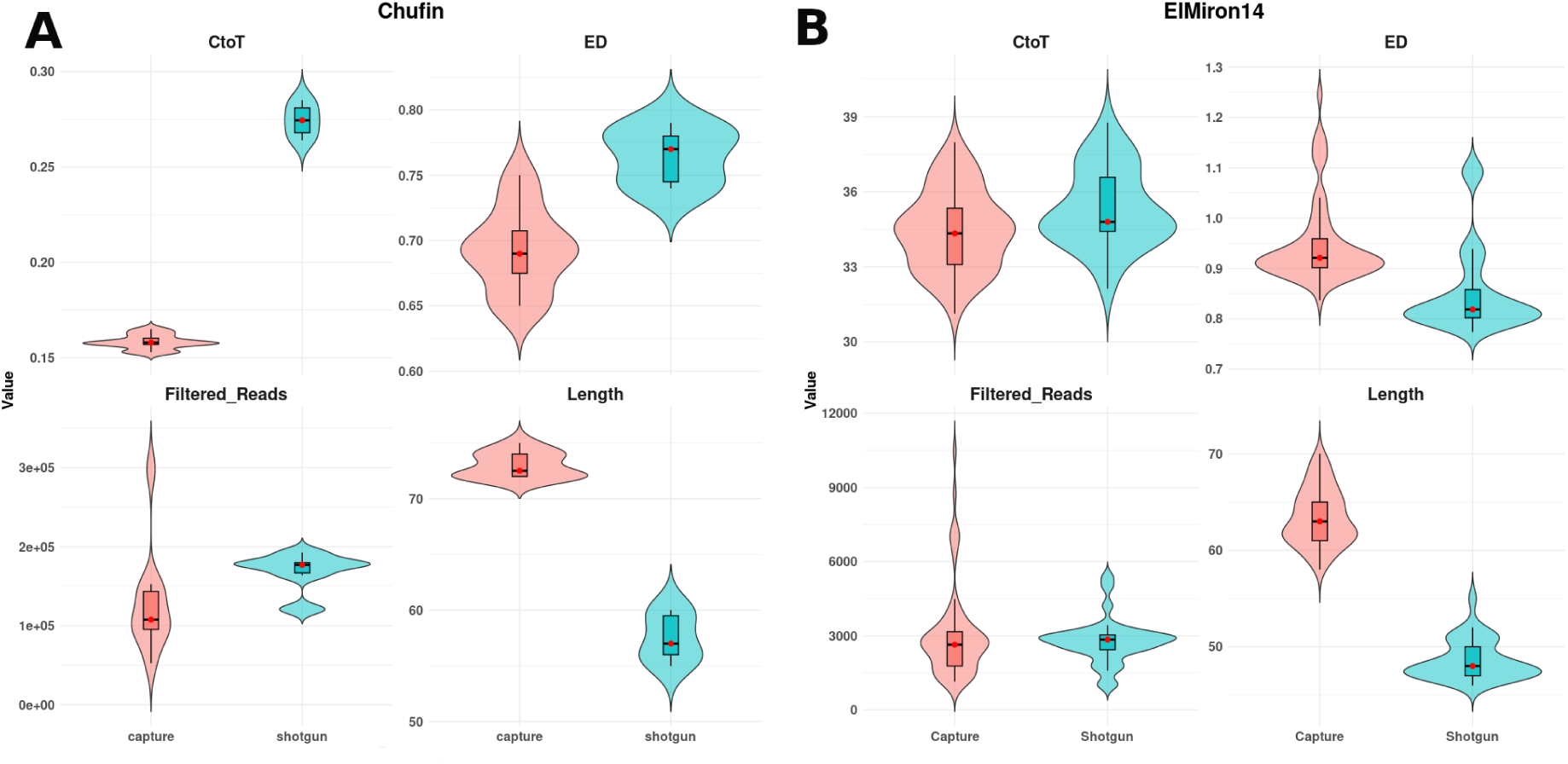
Distribution of key metrics used to characterise the Chufín (A) and ElMiron14 samples (B). The edit distance (ED), deamination frequency at the 5′ end, filtered_Reads (see column U in Data S1), and Length are shown. Red denotes shotgun data, and blue denotes capture data.

Our results, therefore, highlight the significant limitation of using shotgun sequencing for sedaDNA samples when the goal is to recover human genome-wide positions. However, when comparing capture and shotgun libraries, we identified differences across all other parameters analysed, including edit distance (*P=3.392e-08*), read length (*P=4.263e-16*), and deamination in recovered reads (*P=5.483e-16*). The captured reads are, on average, longer and have less deamination. These patterns were also observed in four libraries derived from two separate extracts of powder from a female tooth from Chufín Cave (VIE-1682, 22793 ± 116 yr BP, Supplementary Information, Supplementary Figure 3), which required 52 sequencing attempts (Supplementary Information, Figure 1A). Although some of this could be attributed to modern contamination (Supplementary Data 3, Supplementary Information), the same pattern is observed in samples with no detectable/negligible contamination, such as Ostuni1b, which also indicates that capture reads present longer sequences and less deamination (Supplementary Data 4, Supplementary Information).

Based on the previous evidence and the mtDNA contamination estimates for Chufín (Supplementary Data 3), we determined that the Chufín sample contains between 5% and 8% modern contamination. Although the contamination estimate for ElMiron14 remains undetermined, it also includes modern reads (see Supplementary Information, Supplementary Data 2). Consequently, both individuals were analysed using damage-restricted reads ^17^. In total, we recovered 26,614 SNPs covering the core set of 1,233,013 million SNPs (1240k) targeted both by the Agilent 1240k capture reagent and by the Twist 1.4M capture region for Chufín (‘Chufin_pmd’). By using a combined strategy of shotgun sequencing and in-solution capture, we recovered more than 294,109 unique human reads from the ElMiron14 sediment sample that covered 51,081 SNPs from the 1240k panel ^15^. After damage restriction, 16,239 SNPs from ElMiron14 were retained for analysis (’ElMiron14_pmd’). In addition, we report 467,368 SNPs for Ostuni1b (Supplementary Data 4) (Gravettian male, Italy) and 220,325 SNPs for the Gravettian sample from Isturitz Cave female (southwestern France, 30,850–29,970 cal BP), both with contamination estimates below 5% (Supplementary Data 4).

### East-to-West Gene Flow Highlights Long-Distance Admixture in Southwestern Europe

We analysed the data from the three Gravettian individuals and ElMiron14 Solutrean sedaDNA data, together with the available data from other Upper Palaeolithic individuals, to reconstruct the temporal dynamics of genetic ancestry changes in Western Europe, focusing on the Franco-Cantabrian region. We successfully determined the mtDNA haplogroups of all individuals; all are typical of Upper Palaeolithic and early Holocene individuals in Eurasia ^5,6,18^. Chufín has the mitochondrial U8 haplogroup, which is the same as Dolni Vêstonice 13 ^19^ (Gravettian, Czechia), Goyet Q116-1^5^ (Aurignacian, Belgium) and BK-1653 from Bacho Kiro Cave (Initial Upper Palaeolithic, Bulgaria) ^20^. In addition, a Solutrean individual from Piage (France) ^6^ has the mitochondrial haplogroup U8a, a rare subclade of U8 in present-day Europe but with notably high frequencies among present-day Basques ^21^. Chufín Cave in Cantabria is relatively close to the Basque Country. Isturitz and Ostuni1b carry the M mtDNA haplogroup (as Ostuni1b’s mother, Ostuni1^5^). Haplogroup M is a common haplogroup in Europe before the LGM, but very rare among post-LGM individuals^6,22^. ElMiron14 sediment sample was previously reported to have U2’3’4’7’8’9, a haplogroup observed during the Solutrean with higher frequencies in Western Europe^4,6^. The new sequencing data (Supplementary Information, Supplementary Data 2) support the attribution showing variation within haplogroup U, compatible with multiple individuals.

Using F4ratio^23^, we determined that the genome of Isturitz had about 1.9% Neanderthal ancestry. Ostuni1b had about 1.3%, and the mother (Ostuni1) had 1.5%. These are typical values for the period, and not significantly different from each other ^24,25^. No determination was possible for ElMiron14 or Chufín.

We compared the newly generated genome-wide data from Southwestern Europe (Isturitz, ElMiron14, and Chufín) with the available data from pre-LGM and post-LGM Europe ^4–6^, especially from Western Europe. The Fournol genome from SW France is the oldest available Upper Palaeolithic sequence for SW Europe (29,020-28,450 cal BP). Its ancestry indicates continuity from the Aurignacian-associated individual (represented by Goyet Q116-1(35,630–34,720 cal BP)) to the peak of LGM when the Solutrean culture prevailed in Southern France and Iberia ^26–30^, represented by the Solutrean-associated individual from Malalmuerzo cave in southern Spain (23,020–22,630 cal BP) ^4^. This ancestry would eventually merge with that of Epigravettian-associated individuals from the Italian Peninsula, giving rise to the Magdalenian-associated ancestry in northern Iberia, as represented by the “Red Lady” burial from El Mirón (18,850–18,800 cal BP)^5^.

The three newly reported SW European samples from this study cluster with the previously described Fournol-associated individuals both in an MDS and PCA (Figure 2; Extended Figures 1 and 2). Using several combinations of *f4*(Mbuti, ClusterA; ClusterB, X), we validated that Isturitz, ElMiron14, and Chufín are closer to the Fournol cluster than the Vêstonice cluster (Supplementary Data 5 and 6). These results suggest that the GoyetQ116-1-related ancestry was already present in the Franco-Cantabrian region before 30,000 BP and persisted until approximately 21,000 BP, as is demonstrated by the sedaDNA from the late Solutrean level 122 of El Mirón ^31^ as well as the published data from La Riera (21,010–20,725 cal BP). The newly generated data from the ElMiron14 sample, obtained from sediments and featuring higher SNP coverage than the La Riera sample (Asturias, Spain), supports genetic continuity across the LGM, consistent with human presence in the area linked to the Solutrean technocomplex in the region. The statistic *f*₄*(Isturitz, Goyet Q116- 1; ElMiron14, Mbuti.DG)* yields a non-significant excess of shared genetic drift (z-score < |3|) between ElMiron14 and either Isturitz or Goyet Q116-1. However, due to the low coverage of the ElMiron14 genome, this result does not allow for confident inferences about local population substructure or affinities.

**Figure 2:**
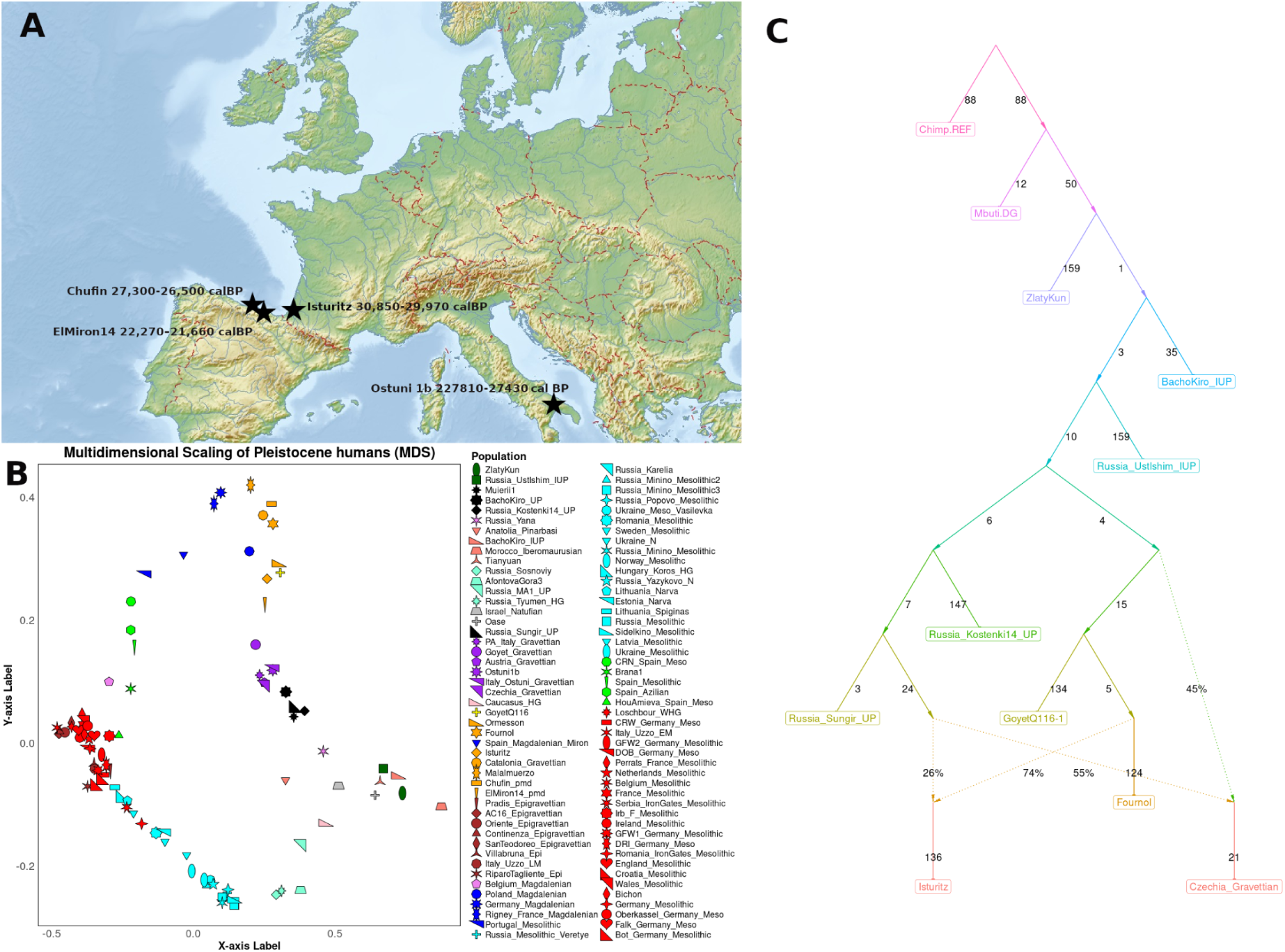
Genomic data from four new individuals from Gravettian-Solutrean associated cultures. A) Location of the sampled individuals. The map was obtained from https://mapswire.com under a CC-BY 4.0 license. B) Chufín, ElMiron14, and Isturitz are associated with the Fournol cluster, Ostuni1b is associated with the Vêstonice cluster, and is closely related to its mother labelled as Ostuni. C) Tree representing the position of Isturitz’s genome based on previous qpGraph analyses from ^6^. The position of Isturitz indicates an influx from a lineage associated with the ancestor of the Sungir genomes

Next, we explored genetic diversity among the Western European LGM genomes using the statistic *f*₄(Mbuti, X; Isturitz, Fournol) (Figure 3B, Supplementary Data 5). The results indicate that all Iberian samples exhibit greater affinity to Fournol than to Isturitz, even though Isturitz is geographically closer to the Iberian region. This pattern may reflect a temporal signal during the Gravettian, as the Isturitz sequence is about 1,000 years older than Fournol or an older ancestry contribution that gets diluted through time. Applying the test *f*₄(Mbuti, X; Isturitz, GoyetQ116-1) (Figure 3A), we found that Věstonice-like individuals exhibit greater affinity to Isturitz than to GoyetQ116-1. In contrast, Solutrean-associated individuals from Iberia are genetically closer to GoyetQ116-1 and Fournol (Figures 3A and 3B). These results suggest that the Solutrean individuals in Iberia may have descended from a Goyet Q116-related population lacking Věstonice ancestry, while a related ancestry could have contributed only to Isturitz. Furthermore, the data raises the possibility that the Isturitz individual received gene flow from eastern or central European populations, which is uncommon in the rest of the southwestern European Palaeolithic individuals of the dataset.

**Figure 3.**
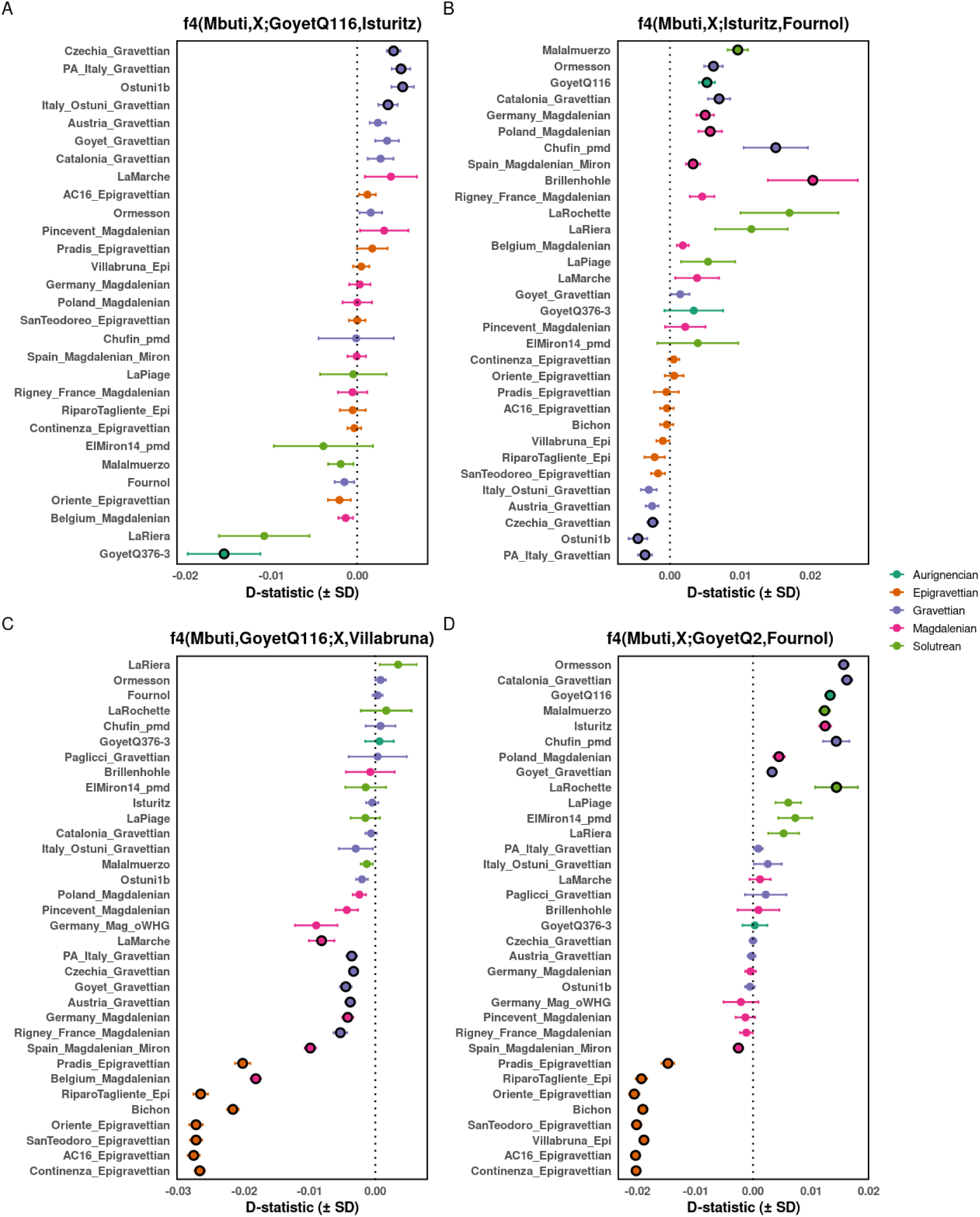
Genomic asymmetries of Isturitz compared to LGM populations of Europe. f4 plots. Empty circles denote results that are not statistically significant (–3 < Z < 3). Colors indicate cultural attribution. Error bars represent one standard error. (A) Gravettian-Věstonice like populations share more derived alleles with Isturitz in comparison to GoyetQ116-1. (B) Western-European Gravettian and Solutrean individuals share more derived alleles with Fournol than with Isturitz. (C) Only the Gravettian-Věstonice like populations and the Magdalenian individuals show clear affinity to Villabruna individual in comparison to GoyetQ116-1 Aurignencian individual. (D) Pre-LGM populations from the Franco-Cantabrian region, including Isturitz, show stronger affinity to Fournol over GoyetQ2.

To explore models that could explain the quantitative values of the *f-statistics* relating the individuals, we fitted admixture graphs using the *find_graphs* function of ADMIXTOOLS 2 ^32^. We used a combination of populations used in Posth et. al 2023^6^, to which we added the genome of Isturitz in 100 independent random searches, producing over 100,000 possible trees (Supplementary Information). The best-fitting graph shows that the genome of Isturitz is consistent with deriving from an admixture of a dominant Fournol-like ancestry with an influx from a population related most closely to the ancestors the individuals labelled Russia Sungir_UP (Russia, 33-31 ka cal BP), from the same lineage Posth et al 2023 previously suggested contributed to Věstonice^5,33^ (Figure 2C, Supplementary Information). The genetic results could be explained by movements of people from Eastern Europe during the Gravettian period, which eventually resulted in this ancestry contribution in Isturitz or by a pre-Goyet Q116 ancestry. The site of Isturitz has long been recognised as a major aggregation (or “super”) site, ^34–41^. So-called aggregation sites refer (albeit with debate) to locations (e.g., large, favourably situated caves) characterised by prolonged and intensive occupation by multiple human bands for a wide variety of purposes, including those of collective hunts, goods and information exchange, socialisation and mating. They are characterised in the Upper Palaeolithic by the presence of abundant portable artworks, often together with rock art, and other archaeological features such as structured living spaces with numerous hearths, large amounts of faunal remains, high artefact density and diversity, and evidence of repeated visits.

To place these results in context, we extended the analysis to Western European individuals associated with the Fournol cluster with >50,000 SNPs (Malalmuerzo, Catalonia_Gravettian and Ormesson; Supplementary Figures 5–7). Surprisingly, we also identified the presence of admixed ancestry in the genome of Ormesson (NW France, 31,822–29,970 cal BP) (Supplementary Information, Supplementary Figure 5), for which Kostenki14 was the best-fitting source, but not in the other tested individuals (Supplementary Figures 6 and 7).

To validate these signals, we computed several tests using Russia_Sungir_UP and Russia_Kostenki14_UP ancestries (Extended Figure 3, Supplementary Data 7). The symmetry statistic *f4*(Mbuti.DG, Russia_Sungir_UP; Fournol, Isturitz)=0.0026, SE=7.52×10-4, Z=3.516, which supports the existence of gene-flow between Isturitz and Eastern populations (after 35,000 BP) when compared to Fournol. This gene-flow is also observed in some Věstonice-like individuals, but not in any published individual from southwestern Europe (Extended Figure 3D). However, the statistic *f4*(Mbuti.DG, Russia_Kostenki14_UP; Fournol, Isturitz)=5.52×10-4, SE=9.18×10-4, Z=0.60, is non-significant (Extended Figure 3B). These same tests are less significant when Goyet-Q116 is placed instead of Fournol (Extended Figure 3A, and 3C). Coincident with the ADMIXTOOLS 2 results, the genome from Ormesson also shows increased affinity to Russia_Kostenki14_UP, but without statistical significance over Goyet-Q116 or Fournol (Extended Figure 3A and B, Supplementary Data 7).

The statistic *f4*(Mbuti.DG, Isturitz; Russia_Sungir_UP, Czechia_Gravettian)=2.931×10^-3^, SE=6.32×10^-4^, Z=4.637 indicates that Isturitz has a greater affinity to Věstonice than to Russia_Sungir_UP, suggesting either gene flow from a population more closely related to Věstonice or an effect of shared ancestry between Věstonice and Goyet-Q116-1 biasing Isturitz’s affinity toward Věstonice. Technical differences in data generation, such as shotgun sequencing versus 1240k capture, can produce spurious affinities unrelated to true population history. For instance, the apparent attraction between Isturitz and Czechia_Gravettian (1240k capture) over Sungir genomes (Shotgun) may reflect shared 1240k ascertainment rather than genuine genetic relationships. A group of Goyet Gravettian-associated individuals from Goyet cave (NW Europe) has been previously reported to be an intermediate between the Fournol and Věstonice clusters ^6^. Here we demonstrate that these individuals are significantly more closely related to Isturitz than to Fournol using *f4*(Mbuti.DG, Goyet_Gravettian; Isturitz, Fournol)=0.0024, SE=7.66×10^-4^, Z=-3.195. Taken together, these results indicate that the Franco-Cantabrian population between 30,000 and 22,000 BP was largely stable, with the notable exception of Isturitz, which shows admixture not observed elsewhere in the region but present in another Western European genome. This highlights the particularity of Isturitz within the broader regional context.

Finally, to determine the level of continuity in El Mirón cave from the Solutrean to the Magdalenian, we use the statistic *f4*(Mbuti, X; Spain_Magdalenian_Miron, ElMiron14). Despite the low coverage, most samples show values close to 0, indicating that only Villabruna-ancestry-related samples exhibit an apparent affinity to Magdalenian ancestry (Figure 3C, Supplementary Figure 8). We then applied the statistic *f4*(Mbuti, X; GoyetQ2, Fournol), which showed that none of the samples from the Cantabrian region associated with the Solutrean or Gravettian show affinity to the GoyetQ2 Magdalenian cluster (Figure 3D). This result confirms the previous findings of the arrival of Villabruna-associated ancestry only after the LGM^5^.

### Inferring the characteristics of Ostuni’s 1b father’s ancestry

Ostuni1b individual is located very close to his mother Ostuni1 in the MDS plot (Figure 2), as expected for a 1st-degree relative. We then tested the possible ancestry of Ostuni1b’s father by comparing Ostuni1b with the mother, Ostuni1, with the symmetry statistic *f4(Mbuti, X; Ostuni1, Ostuni1b).* Any substantial difference in the father’s and mother’s ancestry would move the results in a specific direction. We do not observe any significant differences, so our results are consistent with Ostuni1b’s father and mother being from the same population to the limits of our resolution. Using hapROH^42^ we identified that Ostuni1b had 80 cM in RoH>4cM cM. We did not identify any RoH longer than 8 cM, which indicates a limited population size but no inbreeding. (Extended Figure 4). We successfully classified the Y-chromosome haplogroup of the individual within the C diversity (C*(xC-M217, C-M8, C-V86, C-Y28069, C-B65, C-M208)).

## Discussion and Conclusion

The sequencing of sedaDNA from archaeological contexts has recently enabled the retrieval of genetic material in the absence of osseous remains, widening the possibility to unravel unseen genetic dynamics^7,8,10,13^. However, sedaDNA analyses in archaeological settings are also accompanied by specific challenges, including its contextualisation ^12^, the mixing of genetic data from several individuals, and maximising DNA retrieval. Here, we have presented data illustrating the challenge of retrieving enough endogenous DNA from Pleistocene sediments for high-powered population genetic analyses. Through a systematic comparison between shotgun sequencing and targeted in-solution capture approaches, we demonstrated that only targeted capture is efficient enough to provide useful results. However, this method introduces a preference for longer reads, reduced terminal deamination, and a tendency to capture contaminants over endogenous DNA^43,44^. In our study, shotgun sequencing alone proved insufficient to generate enough informative SNPs for inclusion in population genetic analyses. For example, to obtain just 1,515 SNPs from the Solutrean individual from El Mirón, we had to sequence over one billion reads. Thus, for El Mirón Cave sediments, each informative SNP required an average of 680,000 sequenced reads, underscoring the inefficiency of shotgun approaches for sedaDNA. We could not investigate possible capture biases in more heavily degraded samples, such as the Chufín tooth, due to limited data recovery but we observed that the newly generated Ostuni1b Gravettian individual also shows the presence of bias without contamination. Nevertheless, the observed differences in SNP retrieval from sedimentary samples reinforce the conclusion that targeted capture is currently the only viable method for retrieving informative ancient DNA from sediments.

Through newly generated genomic data from three Upper Palaeolithic individuals from the Franco-Cantabrian region, we identify a previously undocumented Eastern European genetic influence in Western Europe. The genome of a female from Isturitz Cave, dated to approximately 30,000 years ago, reveals Sungir-related ancestry, a signal not previously observed in this region. A similar signal can be discerned in the Ormesson genome, while earlier work identified Věstonice-related ancestry in Gravettian individuals from Goyet (28,000–26,000 BP), who occupy an intermediate genetic position between Věstonice and Fournol ^6^. The parsimonious explanation for this was a cline of Věstonice-Fournol admixture associated with cultural contacts among genetically differentiated and non-homogeneous groups. Our results challenge this parsimonious model: Sungir-related ancestry in Isturitz cannot be explained by derivation from the already admixed Věstonice cluster. Instead, it points to a broader Sungir-like genetic influence affecting both central and southwestern Europe before the establishment of the Věstonice–Fournol cline.

These findings also negate the previously plausible hypothesis that the Franco-Cantabrian region was solely associated with Fournol-related ancestry around 30,000 BP. At the same time, the genetic data from Isturitz support a substantial degree of continuity with other Gravettian samples, reinforcing previous observations of broad, shared genetic patterns during the Gravettian. The presence of Eastern Gravettian ancestry at Isturitz points to long-distance gene flow and positions the site as a point of contact for populations across the continent. Given Isturitz’s exceptional assemblages of portable art objects, among the richest in Southern Europe and with close similarities to items throughout the Pyrenean and Cantabrian regions during different Last Glacial cultural periods ^35–37^, this evidence reinforces the archaeological theory that the site was a key hub of cultural and biological exchange during much of the Upper Palaeolithic, located as it is at the critical passage between continental Europe and the Iberian Peninsula at the western end of the Pyrenees. An alternative explanation is that the Sungir-related signal represents an older ancestry component that persisted in parts of Europe and was subsequently diluted. This would be consistent with its presence in pre-30,000-year-old genomes such as Ormesson but its absence in later Fournol-associated samples. Current data cannot distinguish between these hypotheses, but they collectively indicate that population structure and gene flow during the Gravettian were more complex than previously appreciated.

## Methods

A description of the materials and contexts is presented in the Supplementary Information. ElMiron14, Chufín, and Ostuni1b were processed entirely at the Palaeogenomics Laboratory of the University of Vienna, in dedicated clean-room facilities, following strict contamination control protocols. The Isturitz sample was processed jointly at the University of Vienna and Harvard Medical School, using coordinated procedures. We applied contamination control measures to mitigate the effect of modern DNA contamination. We included negative controls at each step of the wet lab pipeline to control for potential contamination of reagents. 50 mg of sediment or bone was extracted per sample following the protocol from Dabney^45^ with adaptations from Korlević ^46^. Sample ElMiron14 was extracted twice from two different 50 mg aliquots to increase the chances of retrieving human DNA. The DNA was eluted in 50 µL TET buffer (10 mM Tris-HCl, 1 mM EDTA, 0.05% Tween 20, pH 8.0) and double-stranded libraries of 25 µL of the extract, as described in Meyer and Kircher ^47^, were prepared without shearing the DNA into smaller fragments. We used a MinElute PCR Purification kit from Qiagen to clean up the samples, instead of SPRI beads, and eluted in 40 µL of EBT buffer (1 mM EDTA, 0.05% Tween-20). A positive control was added to each library batch using 24 µL of deionised water and 1 µL of a 1:250 dilution of CL104. The number of PCR cycles per sample was determined through real-time PCR. Libraries were double-indexed^48^ and amplified with PfuTurbo Cx HotStart DNA Polymerase from Agilent. A subsequent clean-up was performed with 1.2x NGS clean-up magnetic beads per sample, introducing a size selection. We eluted in 25 µL EBT buffer (1 mM EDTA, 0.05% Tween-20). Each indexed library was amplified further in preparation for in-solution capture using KAPA HiFi HotStart DNA Polymerase and IS5/IS6 as primer pairs ^47^. We enriched some of the libraries (Supplementary Data 1, 2, 3) with the 1.4 million reads from TWIST. Following 16 hours of hybridisation at 65°C and four rounds of washing, we mobilised the target DNA from the probes in a PCR cycler at 95°C for 5 minutes. Another qPCR was performed before amplifying half of the captured library using KAPA HiFi HotStart DNA Polymerase and the primer pairs IS5/IS6. We cleaned up with magnetic beads. Quality controls were performed using Qubit and TapeStation. The captured libraries were sequenced at the Vienna BioCenter Core Facilities (VBCF) on an Illumina NovaSeq 6000 platform.

### Alignment of genomic data

Sequencing reads were trimmed using cutadapt (version 1.2.1)^49^ with a minimum length of 30 and removal of polyA tails. We also processed the sequencing reads using fastp 0.25.0 ^50^ with the following parameters (--trim_poly_g --trim_poly_x -poly_x_min_len 5 -poly_g_min_len 5 --length_required 30 --qualified_quality_phred 30). This pipeline ensured the retention of high-quality reads. The resulting reads were aligned to the human reference genome (GRCh37) and the revised Cambridge reference sequence (rCRS) mitochondrial genome using BWA aln^51^ (Version 0.7.5), with seeding disabled and a gap penalty of 2 set for the open end. Duplicate mapped reads were removed using Picard Tools 2.21.4^52^ with default parameters. Reads with mapping qualities below 30 were also removed. Unique and filtered reads were analysed with Qualimap-2.2.1^53^ to assess whole-genome coverage depths. MapDamage-2.0.9 ^54^ was used to estimate the level of deamination and the authenticity of the data.

### Processing of ElMiron14 data

For ElMiron14 reads, we converted the resulting BAM files into FASTA files using Samtools 1.12 ^55^. These sequences were processed with BLASTn 2.16.0 ^56^ using the whole nucleotide database from NCBI (2024-06-07). The results were analysed in MEGAN 6.25.10^57^ using the following parameters: Native LCA, bitscore 50, minimum identity of 80, and top score 10, grouping the Hominidae reads. The resulting reads were subtracted from the original BAM file and realigned against the human reference using the same parameters.

For the mitochondrial DNA (Supplementary Table 14), we used the same methodological approach described in Gelabert et al, 2025, ^7^, capturing with commercial myBaits mtDNA baits. These libraries were sequenced in paired-end 150 reaction reads on a Novaseq X platform, Vienna BioCenter Core Facilities (VBCF). The reads were then clipped using cutadapt (version 1.2.1) with a minimum length of 30 and removal of polyA tails. The clipped reads were then collapsed with PEAR 0.9.11^58^ with a minimum overlap of 11 bases. The resulting reads were aligned against the rCRS mtDNA reference genome using BWA aln^51^ (Version 0.7.5), with seeding disabled and a gap penalty of 2 set for the open end. Duplicate mapped reads were removed using Picard Tools 2.21.4^52^ with default parameters. Reads with mapping qualities below 30 were also removed. Unique and filtered reads were analysed with Qualimap-2.2.1^53^ to assess whole-genome coverage depths. MapDamage-2.0.9 ^54^ was used to estimate the level of deamination and the authenticity of the data. The reads were then converted to fastq using Samtools 1.12^55^. These sequences were processed with BLASTn 2.16.0 ^56^ using the whole nucleotide database from NCBI (2024-06-07). The results were analysed in MEGAN 6.25.10^57^ using the following parameters: Native LCA, bitscore 50, minimum identity of 80, and top score 5, grouping the Primate reads. These were aligned again to the human reference genome and processed with Schmutzi 2.0.0 ^59^ to determine the consensus sequence. We finally examined each of the positions with Samtools mpileup and called the most abundant haplogroup with Haplogrep 3.2.1^60^. We plotted the deamination and read length distributions.

### Quality control and post-processing

To determine the presence of contamination, we applied three methods based on mtDNA diversity: Schmutzi 2.0.0 ^59^, ContamMix^5^, and Calico 2.0^61^. In the case of Ostuni1b, we also used angsd 0.941^62^ to determine contamination in the X chromosome. After this determination of contamination, we processed the reads of Chufin and ElMrion14 with pmdtools 0.60 ^17^, keeping reads with PMD scores greater than 3. Finally, we used GATK 3.8.1.0^63^ to realign indels using RealignerTargetCreator and IndelRealigner, along with Mills_and_1000G_gold_standard.indels.b37.vcf database. We then determined deamination using MapDamage 2.2.2 ^54^, and we trimmed the reads accordingly using bamutil-1.0.15 trimbam option ^64^, removing bases until the last base had <5% of damage. Genotypes were called using the pileup caller of sequenceTools ^65^, filtering out bases with a base-quality score below 30. At this stage, we called genotypes at the positions of the 1240k SNP targets. These calls were merged with ancient and present-day individuals. ^4–6,15,18,20,33,61,66–91^ using PLINK 1.9 ^92^. The full list of samples is available in Data S1D.

We generated consensus sequences of the mtDNA genomes using Schmutzi 2.0.0 and predicted the mtDNA haplogroup using Haplogrep 3.2.1 ^60^. We predicted the Y-chromosome of the Ostuni1b individual using Yleaf v3.2.1^93^, restricting the analyses to positions with three reads covered.

### Population genetics analyses

We used 1074 individuals from Western Eurasian populations from the AADR repository^16^ genotyped on the 597,573 positions of the Affymetrix Human Origins SNP array to perform Principal Components Analysis with smartPCA ^23^. In this PCA, we projected all the samples we report in this paper and other relevant Pleistocene samples (Supplementary Data 3). We used the same dataset to perform *f-statistics*-based analyses using AdmixTools 8.0.2. (Supplementary Information), following standard protocols in paleogenomics. The results were plotted using R 4.4.2^94^. We constructed an *f_3_* matrix using all samples with more than 10,000 SNPs and grouped the samples based on former publications. The results were processed with R to build an MDS plot.

We performed an admixture graph search using the find_graph() function from ADMIXTOOLS 2.0.2 ^32^. To broadly sample topology space and reduce the risk of convergence to local optima, we performed 20 independent searches, each allowing up to two admixture events (max_admix = 2), rooting the graph with Pan troglodytes (“Chimp.REF”), and using stop_gen = 50 generations with numgraphs = 50 candidate graphs per generation. Each run was initialized with a different random seed and parallelized across all available CPU cores. All results were combined, and model fit was ranked using the score returned by find_graphs() (negative log-likelihood, with lower values indicating better fit). The best-fitting topology was defined as the single graph with the lowest score across all runs, and we quantified how often this topology recurred among all candidate graphs. The best model and all results were saved for reproducibility, and we visualized the winning topology. The initial tree structure was informed by the model presented in Posth et al 2023 ^6^.

We applied the ROH analysis approach by ^42^, but only to the Ostuni1b genome. We plotted the results with the Python scripts at (https://github.com/hringbauer/hapROH).

## Materials availability

The remaining material from the analysed samples is available and stored under secure conditions at the University of Vienna for further research, subject to an agreement with the research team.

## Data and code availability

Sequencing reads are available in the European Nucleotide Archive (ENA) under accession number PRJEB90506.

## Acknowledgements

This research was funded by the European Research Council (ERC) under the SHADOWS project (grant agreement No. 101163057) and by the INEAL STMG grant CA19141-8d068698 (P.G.). J.-M.T. is supported by a project of the Meitner Programme of the Austrian Science Fund (FWF) (Project: *Osseous Hunting Weapons of Early Modern Humans in Eurasia*, No. M3112), by the Ramón y Cajal programme of the Spanish Ministry of Science and Innovation (MCIN/AEI/10.13039/501100011033, grant RYC2021-033759-I), and by the European Union (NextGenerationEU/PRTR). V.V.-M. is supported by a ‘Ramón y Cajal’ contract (grant RYC2022-035700-I), funded by the Ministerio de Ciencia, Innovación y Universidades. D. CS was supported by project PID2021-124589NA-I00, funded by MCIN/AEI/10.13039/ 501100011033 and the European Union (FEDER) and from the Ramón y Cajal program (grant RYC2022-036585-I), funded by the Spanish Ministry of Science and Innovation. A.B. M.-A. was supported by the European Research Council (ERC) under the SUBSILIENCE project (grant agreement No. 818299). A.N. is supported by the European Research Council (ERC) under the European Union’s Horizon Europe Research and Innovation Programme (project MOTHERS, grant agreement no. 101077348). M. S. and S. S. are supported by the European Research Council (ERC) under the MicroStratDNA project (grant agreement 101042570). The Research Platform MINERVA also supported the research at the University of Vienna. Fieldwork and research at El Mirón Cave received funding from the Dirección General de Cultura y Patrimonio Histórico de la Consejería de Cultura, Turismo y Deporte del Gobierno de Cantabria, the US National Science Foundation, Fundación M. Botín, the Leakey Foundation, the Ministerio de Educación y Ciencia, the National Geographic Society, the University of New Mexico, and the Stone Age Research Fund at the UNM Foundation (J. and R. Auel principal donors). R.P. and D.R. were supported by John Templeton Foundation grant 61220. D.R. was supported by National Institutes of Health grant HG012287 and by the Allen Discovery Centre programme, a Paul G. Allen Frontiers Group-advised programme of the Paul G. Allen Family Foundation, and is an Investigator of the Howard Hughes Medical Institute. Additional material support was provided by the Town of Ramales de la Victoria and the Instituto Internacional de Investigaciones Prehistóricas de Cantabria. Fieldwork and research at Chufín Cave received funding from the Gobierno de Cantabria and Palarq Foundation. We thank the archaeological teams from the University of Cantabria and New Mexico working at El Mirón and Chufín caves for their support and assistance with sampling and contextual documentation. We also gratefully acknowledge the team at the Isturitz and Oxocelhaya caves, as well as the Darricau family, for their support during this study.

## Author Contributions

P. G, R. P, S. S conceptualised the study. L. G. S, MR. GM, I. GZ, D. CS, V. O, JM T, D. G and P. G sampled, V. O, F. E, B. Z, F. B, F. T C, S. F, C. G-M, M. H, JM T, T. H, L. vD S, M. G and S. S performed the experiments, P. G, V.V-M, V. O, M. H, C G-M, and D. R analysed the data. C. N, J. D, L-G. S, M R. G M, I. G Z, D. C S, AB. M A, D. G, A. C, D. C, M. S. provided archaeological material, context, and contributed to the experimental design of the manuscript. P. G, R. P. and D.R wrote the text with input from all collaborators.

---

**Extended Figure 1:**
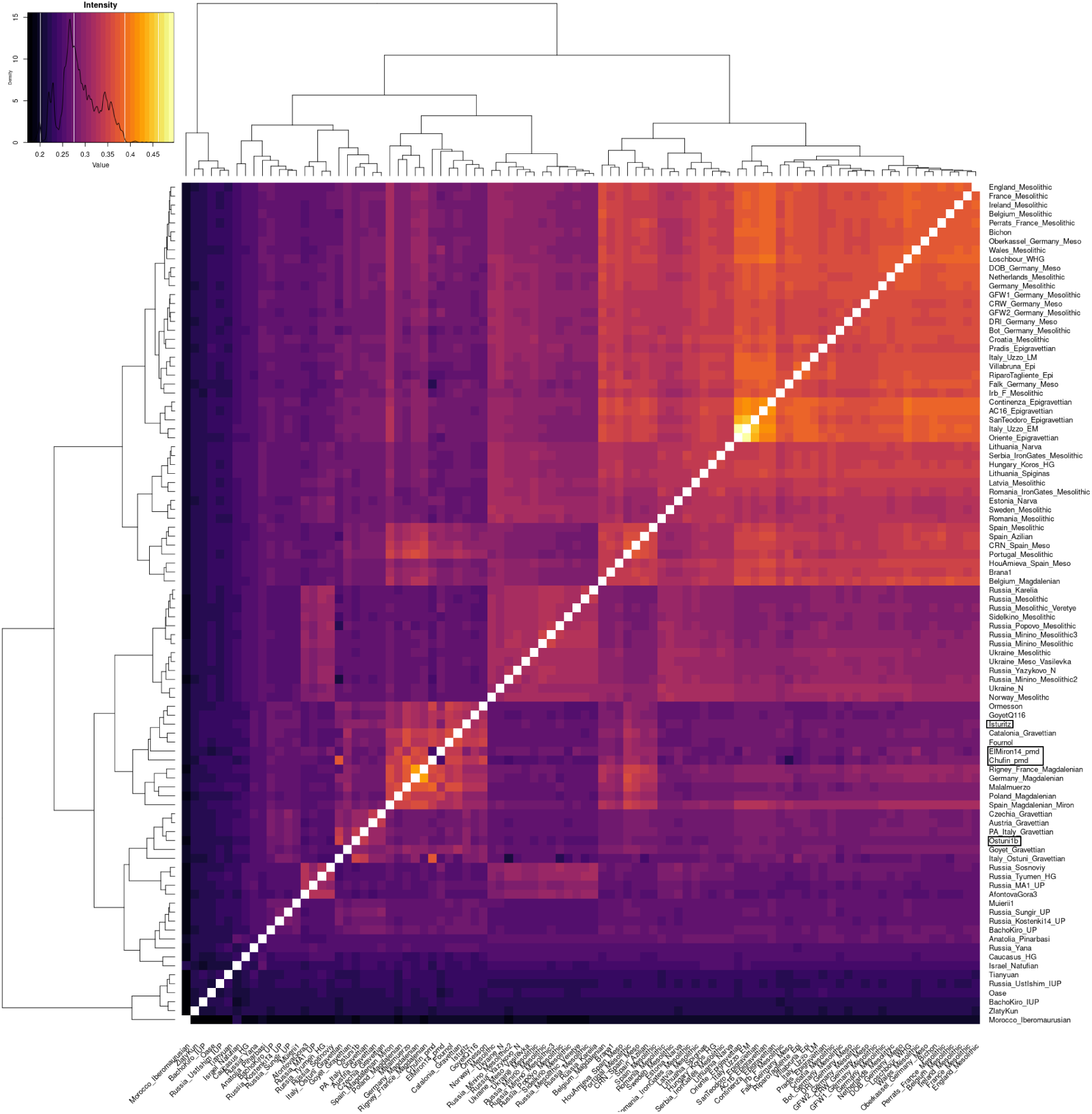
f_3_ matrix reflecting genetic distances among all possible pairs of individuals and calculated as f_3_(HGi, HGj; Mbuti) using all the samples with more than 10,000 SNPs and grouping the samples based on former publications covering the UP populations. The newly generated Isturitz, ELMiron_14 and Chufin are located in the Fournol Cluster. Ostuni1b is almost identical to Ostuni. The four newly reported samples are marked on the y-axis.

**Extended Figure 2:**
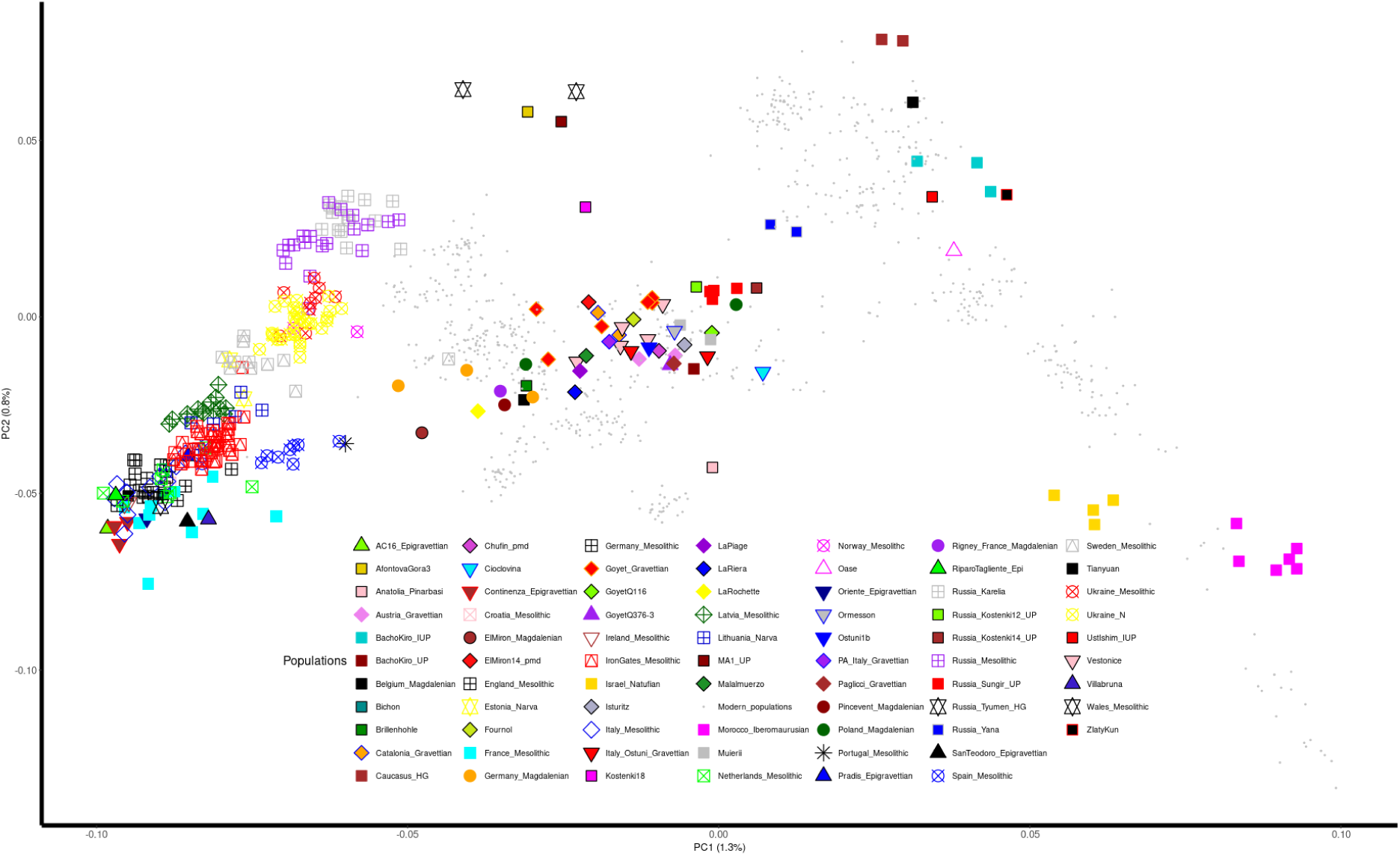
Principal Components analysis performed with European modern genomes in which samples from Supplementary Data 4 have been plotted.

**Extended Figure 3:**
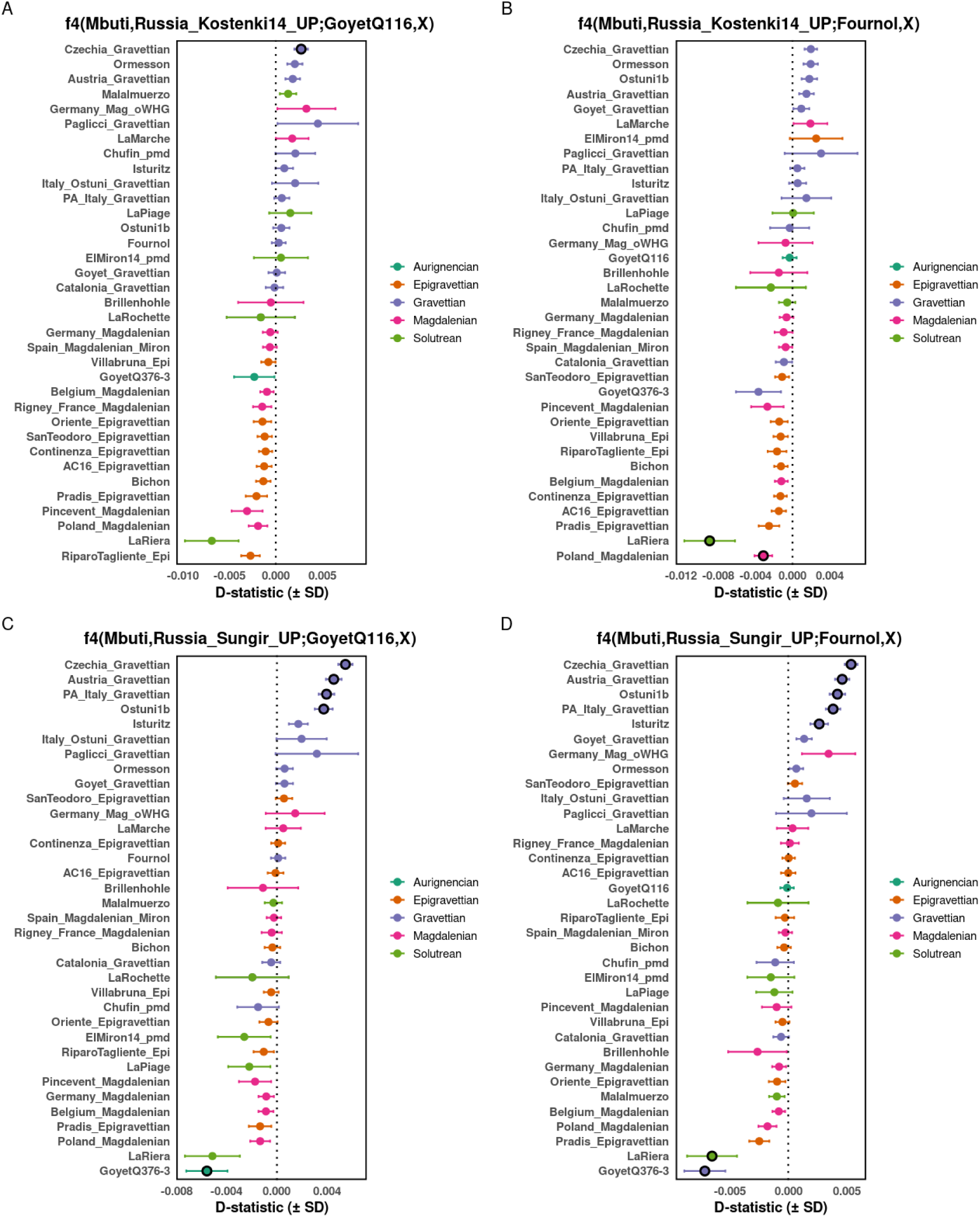
Genomic asymmetries of Upper Pleistocene European populations compared to the Russia_Sungir_UP and Russia_Kostenki14_UP. The plots show that Russia_Kostenki14_UP shows affinity to Chezkia_Gravettian and to Ostuni (A, B). Russia_Sungir_UP shows affinity for Isturitz over Fournol (D), showing a similar relationship to other Eastern Gravettian populations (C). Empty circles denote no statistical significance (-3>Z<3), the colours show cultural attribution. Error bars indicate one standard error.

**Extended Figure 4:**
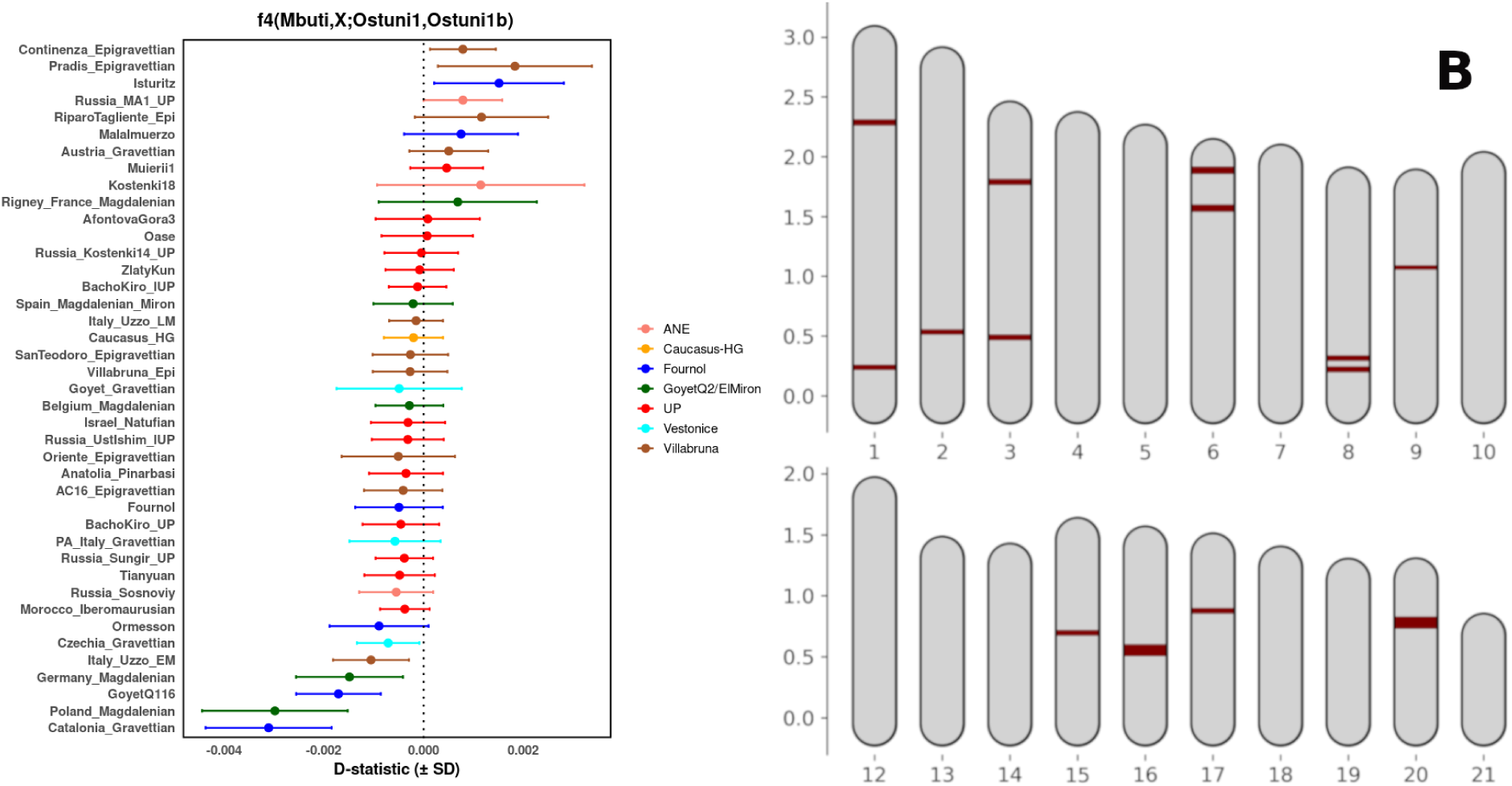
Ostuni1 b: **A)** Comparison of Ostuni1b and Ostuni1 in the form f4(Mbuti, X; Ostuni1, Ostuni1 b). Individuals on the positive side of the plot show a greater affinity to Ostuni1b, while those on the left side are classified by genetic cluster as individuals. B) RoH of Ostuni1b.

