## Supplementary file for "Genetic admixture between East and West European Gravettian-associated populations in Western Europe before the Last Glacial Maximum"

### Supplementary Information

### 1. El Miron sediments

#### 1.1 Sediment Processing from El Mirón Cave

ElMiron14 is a soil sample from square W10 of level 122 in El Mirón Cave, dated to 22,270–21,660 calBP [1], corresponding to the Late Solutrean period in Cantabria. In a previous study where this sample was enriched using mtDNA baits [2], we found that it contained substantial amounts of human DNA, and is the sample with the highest number of human reads from the ones screened to date from El Mirón Cave. To maximise the recovery of human genomic reads, we implemented a multiple-recovery strategy using 101 sequenced libraries deriving from a 100mg soil sample. This included three different extracts and a total of nine double-stranded libraries. These libraries were sequenced using a combination of in-solution baits covering the 1240k positions (manufactured by TWIST Bioscience) and shotgun sequencing. Altogether, 1,028,866,146 reads were generated across all libraries. Of these, 652,257 were classified as Hominin based on BLAST results, and finally, 283,315 of these could be confidently attributed to human ancestry (see Star Methods).

Next, we performed in-solution capture on some of the libraries. The complete list is available in Supplementary Data 1. In summary, we prepared nine libraries from the El ElMiron14 sample. These libraries are: l1980 and l2594 from extract 1987, and libraries 2593, 2592, 2589, 2588, 2236, 2590, and 2591 from extract 2668. These libraries were re-indexed and sequenced a total of 101 times.

#### 1.2 Diversity between the samples

We tested the presence of diversity in the data using four parameters: read deamination, read edit distance, length distribution, and sequencing yields, comparing the libraries and the extracts. We aimed to detect any difference among the 101 sequenced libraries. We computed the mean, median, and SD of these parameters, and we tested the presence of significant differences using the Mann-Whitney U test, all integrated in R 4.4.2 [3]

We first tested for differences in Edit distance (Supplementary Table 1-2) across sequencing attempts organised by libraries and extracts that would potentially suggest that the sequenced material has a different nature.

Supplementary Table 1: Computed statistics on the Edit distance per Extract ID.

| **ExtractID** | **mean** | **median** | **sd** | **min** | **max** |
|---|---|---|---|---|---|
| 1987 | 0.926 | 0.924 | 0.109 | 0.774 | 1.17 |
| 2668 | 0.916 | 0.906 | 0.0928 | 0.784 | 1.25 |

After performing the Mann-Whitney U test (Wilcoxon rank-sum test), the p-value e of 0.469 indicates no statistically significant difference between extracts regarding edit distance.

Supplementary Table 2: Computed statistics on the Edit distance per Library ID.

| **LibraryID** | **mean** | **median** | **sd** | **min** | **max** |
|---|---|---|---|---|---|
| l1980 | 1.00 | 0.949 | 0.147 | 0.888 | 1.17 |
| l2236 | 1.09 | 1.09 | 0.0477 | 1.02 | 1.16 |
| l2588 | 0.930 | 0.932 | 0.0502 | 0.837 | 1.00 |
| l2589 | 0.873 | 0.891 | 0.0499 | 0.784 | 0.929 |
| l2590 | 0.899 | 0.888 | 0.117 | 0.795 | 1.25 |
| l2591 | 0.883 | 0.912 | 0.0648 | 0.790 | 0.954 |
| l2592 | 0.885 | 0.905 | 0.0505 | 0.798 | 0.934 |
| l2593 | 0.875 | 0.884 | 0.0426 | 0.802 | 0.957 |
| l2594 | 0.908 | 0.922 | 0.0968 | 0.774 | 1.11 |

After performing the Kruskal-Wallis chi-squared test = 36.236, df = 8, $P$= 1.59e-05. Despite these differences, as the extracts showed no statistical difference, we do not consider that certain libraries could be problematic.

We next tested the presence of bias in the read lengths, which may suggest differential modern contamination (Supplementary Table 3-4).

Supplementary Table 3: Computed statistics on the read length per Extract ID.

| ExtractID | mean | median | sd | min | max |
|---|---|---|---|---|---|
| 1987 | 61.8 | 65 | 7.68 | 50 | 70 |
| 2668 | 57.9 | 61 | 7.36 | 46 | 69 |

After performing the Mann-Whitney U test (Wilcoxon rank-sum test), the *P*-value is 0.011, indicating a statistical difference (at $P<0.05$) between extracts in terms of read length.

Supplementary Table 4: Computed statistics on the read length per Library ID.

| LibraryID | mean | median | sd | min | max |
|---|---|---|---|---|---|
| l1980 | 64.7 | 69 | 8.39 | 55 | 70 |
| l2236 | 60.5 | 62 | 6.52 | 51 | 69 |
| l2588 | 56.7 | 60 | 7.33 | 47 | 65 |
| l2589 | 58.2 | 61.5 | 7.62 | 48 | 66 |
| l2590 | 57.6 | 60 | 7.95 | 47 | 67 |
| l2591 | 56.4 | 59.5 | 7.29 | 46 | 64 |
| l2592 | 58.6 | 62 | 8.31 | 47 | 68 |
| l2593 | 57.9 | 61 | 7.50 | 47 | 65 |
| l2594 | 61.1 | 64.5 | 7.72 | 50 | 69 |

After performing the Kruskal-Wallis chi-squared test value 12.234, df = 8, $P = 0.141$, we observe no differences in read length across the libraries.

We then tested differences across sequencing libraries and extracts regarding the deamination in the 5' end (Supplementary Tables 5 and 6).

Supplementary Table 5: Computed statistics on the deamination at 5' per extract ID.

| ExtractID | mean | median | sd | min | max |
|---|---|---|---|---|---|
| 1987 | 0.412 | 0.389 | 0.0453 | 0.356 | 0.482 |
| 2668 | 0.410 | 0.389 | 0.0574 | 0.298 | 0.532 |

After performing the Mann-Whitney U test (Wilcoxon rank-sum test), the $P$=0.699 indicates no statistically significant difference between extracts regarding read length.

Supplementary Table 6: Computed statistics on deamination at 5’ per Library ID.

| LibraryID | mean | median | sd | min | max |
|---|---|---|---|---|---|
| l1980 | 0.405 | 0.387 | 0.0597 | 0.356 | 0.471 |
| l2236 | 0.401 | 0.388 | 0.0506 | 0.310 | 0.472 |
| l2588 | 0.421 | 0.389 | 0.0622 | 0.358 | 0.532 |
| l2589 | 0.392 | 0.368 | 0.0641 | 0.298 | 0.503 |
| l2590 | 0.408 | 0.380 | 0.0577 | 0.344 | 0.519 |
| l2591 | 0.437 | 0.422 | 0.0484 | 0.381 | 0.516 |
| l2592 | 0.411 | 0.384 | 0.0613 | 0.347 | 0.507 |
| l2593 | 0.403 | 0.382 | 0.0582 | 0.330 | 0.505 |
| l2594 | 0.414 | 0.394 | 0.0442 | 0.366 | 0.482 |

We tested the presence of bias across libraries regarding damage, and we did not identify it: Kruskal-Wallis chi-squared = 8.7986, df = 8, *P*= 0.360.

We finally tested differences in human-read recovery per sample grouped by extract ID and library ID (Supplementary Tables 7 and 8).

Supplementary Table 7: Computed statistics on human reads recovery per Extract ID.

| LibraryID | mean | median | sd | min | max |
|---|---|---|---|---|---|
| 1987 | 5294 | 5000 | 2154 | 2870 | 10480 |
| 2668 | 2523 | 2612 | 1015 | 993 | 8757 |

After performing the Mann-Whitney U test (Wilcoxon rank-sum test), $P=3.75\times10^{-8}$, showing a statistical difference between extracts regarding human recovery.

Supplementary Table 8: Computed statistics on human reads recovery per library ID.

| LibraryID | mean | median | sd | min | max |
|---|---|---|---|---|---|
| l1980 | 7599 | 6930 | 2612 | 5387 | 10480 |
| l2236 | 3065 | 2557 | 2104 | 1063 | 8757 |
| l2588 | 2334 | 2487 | 626 | 1147 | 3145 |
| l2589 | 2344 | 2622 | 634 | 1183 | 3084 |

| | | | | | |
|---|---|---|---|---|---|
| l2590 | 2450 | 2500 | 791 | 1178 | 3918 |
| l2591 | 2550 | 2623 | 910 | 993 | 4090 |
| l2592 | 2554 | 2820 | 590 | 1573 | 3115 |
| l2593 | 2427 | 2628 | 853 | 1217 | 4213 |
| l2594 | 4717 | 3802 | 1689 | 2870 | 7243 |

We tested the presence of bias across libraries regarding human reads, and we did not identify it: Kruskal-Wallis chi-squared = 8.7986, df = 8, p-value = 0.3596.

After these comparisons, we repeated the same strategy and performed comparisons on ED, length, 5' damage, and human recovered reads, this time classifying the sequencing attempts by capture and shotgun (Supplementary Data 1, Supplementary Table 9-10).

Supplementary Table 9: Computed statistics on human reads recovery per sequencing strategy.

| **Method** | **mean_ed** | **median_ed** | **sd_ed** | **min_ed** | **max_ed** |
|---|---|---|---|---|---|
| Capture | 0.950 | 0.922 | 0.0830 | 0.837 | 1.25 |
| Shotgun | 0.855 | 0.819 | 0.0883 | 0.774 | 1.10 |

We performed a Mann-Whitney U test, and $P=2.86\times10^{-8}$, showing that shotgun sequencing has lower ED than capture-recovered reads.
Next, we tested the presence of differences in length classified by method.

Supplementary Table 10: Computed statistics on human reads length per sequencing strategy.

| **Method** | **mean** | **median** | **sd** | **min** | **max** |
|---|---|---|---|---|---|
| Capture | 63.4 | 63 | 2.92 | 58 | 70 |
| Shotgun | 48.6 | 48 | 2.02 | 46 | 55 |

We performed a Mann-Whitney U test, and the p-value is $4.26\times10^{-16}$ which indicates that capture samples are significantly longer than the shotgun ones.

We then tested the differences in the deamination at the 5’ end of human reads recovered (Supplementary Tables 11 and 12).

Supplementary Table 11: Computed statistics on human read deamination per sequencing strategy.

| **Method** | **mean** | **median** | **sd** | **min** | **max** |
|---|---|---|---|---|---|
| Capture | 0.376 | 0.379 | 0.0260 | 0.298 | 0.442 |
| Shotgun | 0.481 | 0.476 | 0.0221 | 0.449 | 0.532 |

The *P*-value of a Mann-Whitney U test (Wilcoxon rank-sum test) is $5.48\times 10^{-16}$, showing that capture reads have much less deamination.

We then tested the differences in the quantity of human reads recovered.

Supplementary Table 12: Computed statistics on human read recovery per sequencing strategy.

| **Method** | **mean** | **median** | **sd** | **min** | **max** |
|---|---|---|---|---|---|
| Capture | 2976 | 2636 | 1852 | 214 | 10480 |
| Shotgun | 2779 | 2846 | 902 | 993 | 5387 |

The *P*-value of a Mann-Whitney U test (Wilcoxon rank-sum test) is 0.636, showing no differences in recovery of human reads between capture and shotgun libraries.

We then performed several tests to observe and quantify the differences in our merged files: Merged_Shotgun, Merged_Capture, Merged, and Merged_under90bp. We then called pseudo-haploid genotypes using PilupCaller from Sequencetools 1.6.0.0 [4]. The results are presented in Supplementary Table 13.

Following the observations of the deamination values, we clipped five reads per read end using trimbam from bamutil-1.0.15 [5]. The resulting final Merged file has 283,315 reads, an average fragment length of 59bp, and 40% of damage.

Supplementary Table 13 Results from the PileupCaller on the four samples generating merging ElMiron_14 reads in the four combinations presented.

| SampleName | TotalSites | Covered calls |
|---|---|---|
| Miron14 | 1233013 | 49212 |
| Miron14_S | 1233013 | 1515 |
| Miron14_C | 1233013 | 48384 |
| Miron14_90 | 1233013 | 42214 |

We first used *f3*-statistics in the form *f3*(X,Y: Mbuti) to determine the similarity between the different comparisons of the Miron sample.

Supplementary Table 13: Results of the *f3*-statistics computed on ElMiron_14 samples and other contemporary and regional genomes.

| Pop1 | Pop2 | Pop3 | Value | SD | SNPs |
|---|---|---|---|---|---|
| ElMiron14 | ElMiron14_C | Mbuti | 0.786272 | 0.007524 | 30795 |
| ElMiron14 | ElMiron14_90 | Mbuti | 0.783615 | 0.008001 | 26803 |
| ElMiron14 | ElMiron14_S | Mbuti | 0.733932 | 0.034405 | 942 |
| ElMiron14_C | ElMiron14_90 | Mbuti | 0.785554 | 0.008061 | 26318 |
| ElMiron14_90 | ElMiron14_S | Mbuti | 0.714201 | 0.035299 | 866 |
| ElMiron14_C | ElMiron14_S | Mbuti | 0.740460 | 0.054262 | 438 |
| Href.REF | ElMiron14 | Mbuti | 0.236326 | 0.006225 | 32954 |
| Href.REF | ElMiron14_C | Mbuti | 0.235931 | 0.006247 | 32426 |
| Href.REF | ElMiron14_S | Mbuti | 0.248459 | 0.029919 | 985 |
| Href.REF | ElMiron14_90 | Mbuti | 0.241878 | 0.006726 | 28213 |
| Href.REF | Isturitz | Mbuti | 0.193309 | 0.003867 | 153538 |
| Href.REF | GoyetQ116 | Mbuti | 0.190310 | 0.003449 | 411468 |
| Href.REF | GoyetQ2 | Mbuti | 0.199044 | 0.003611 | 293798 |

| | | | | | |
|---|---|---|---|---|---|
| Href.REF | Girona_Grav | Mbuti | 0.241513 | 0.004110 | 136327 |
| Href.REF | ElMiron_Mag | Mbuti | 0.205492 | 0.003555 | 434089 |
| Href.REF | Fournol | Mbuti | 0.212010 | 0.003654 | 408064 |
| Href.REF | Malalmuerzo | Mbuti | 0.215603 | 0.004039 | 151253 |
| Spanish.DG | Isturitz | Mbuti | 0.249433 | 0.003267 | 165272 |
| Spanish.DG | GoyetQ116 | Mbuti | 0.245307 | 0.002884 | 442897 |
| Spanish.DG | GoyetQ2 | Mbuti | 0.256118 | 0.003026 | 315798 |
| Spanish.DG | Girona_Grav | Mbuti | 0.247653 | 0.003308 | 148083 |
| Spanish.DG | ElMiron_Mag | Mbuti | 0.257933 | 0.003003 | 465486 |
| Spanish.DG | Malalmuerzo | Mbuti | 0.251052 | 0.003393 | 163038 |
| Spanish.DG | Fournol | Mbuti | 0.247220 | 0.003028 | 441054 |
| Spanish.DG | ElMiron14 | Mbuti | 0.253243 | 0.004752 | 35711 |
| Spanish.DG | ElMiron14_C | Mbuti | 0.253971 | 0.004791 | 35125 |
| Spanish.DG | ElMiron14_S | Mbuti | 0.237300 | 0.021585 | 1081 |
| Spanish.DG | ElMiron14_90 | Mbuti | 0.251281 | 0.00504 | 30606 |

Without proper contamination estimates due to the multi-individual origin of the ElMiron14 reads, we decided to apply a PMD restriction using PMD tools 0.60 [6]. The process resulted in 95,843 reads. We then trimmed 5 bases per read end. The process resulted in 16,239 SNPs covering the 1240k SNP target set. These highly confident SNPs are the ones we used for the ancestry determination of the ElMiron14_pmd sample.

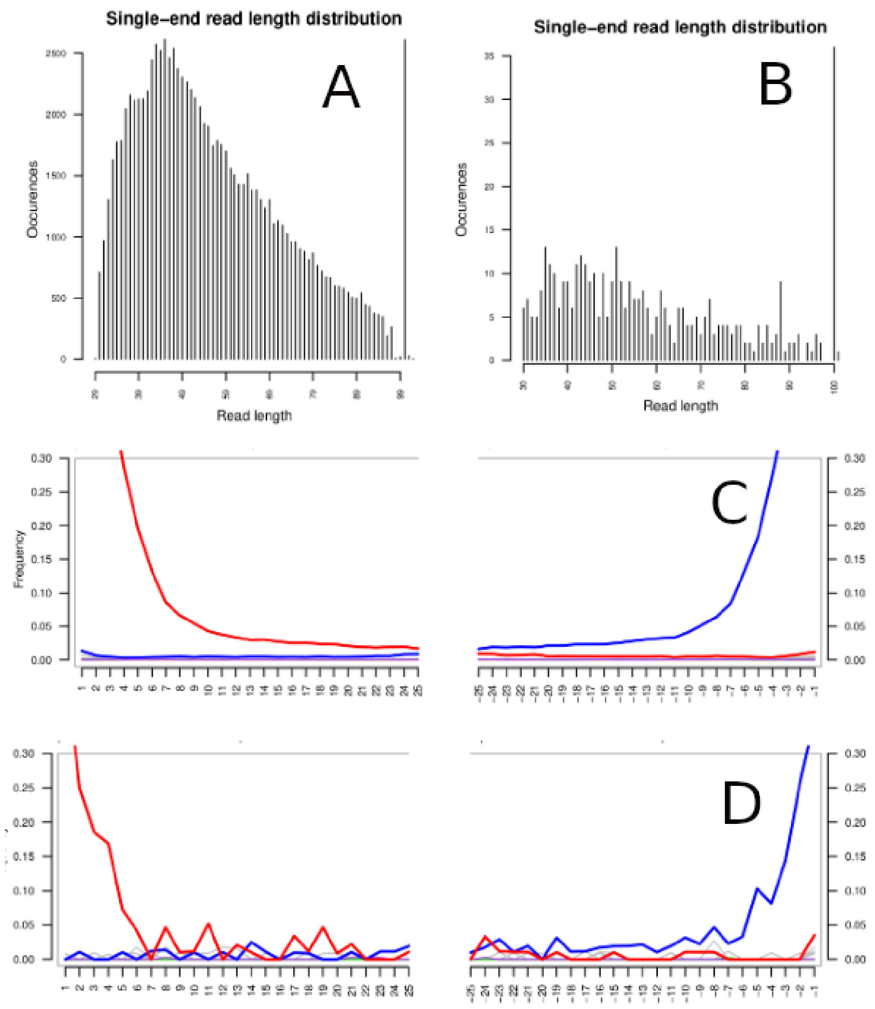


**Supplementary Figure 1: ElMiron14 sequencing.** A) Distribution of read lengths of ElMiron14_pmd, showing characteristic fragmentation patterns typical of degraded ancient DNA, B) Distribution of read lengths of ElMiron14 mtDNA, showing characteristic fragmentation patterns typical of degraded ancient DNA, C) Cytosine deamination patterns along sequencing reads of ElMiron14_pmd, indicative of ancient DNA authenticity. D) Cytosine deamination patterns along sequencing reads of ElMiron14 mtDNA, indicative of ancient DNA authenticity.

#### 1.3 The Mitochondria of Miron 14

We explored the retrieval of mitochondrial DNA from the 101 sequencing attempts of the 9 libraries. After merging all the reads and removing duplicates, we obtained only 418 unique sequences in total. This result indicates that the capture baits used in the experiment did not allow for substantial mtDNA recovery from the shotgun libraries, robably due to an error in the experimental side (Supplementary Figure 1).

We then applied a mitochondrial capture procedure following the same methodological approach described in Gelabert et al, 2025, [2], capturing with commercial myBaits mtDNA baits. The sequenced libraries are presented in Supplementary Table 14. All the libraries show short reads and substantial deamination (Supplementary Figure 2). The results of the calls are presented in Supplementary Data 2.

**Supplementary Table 14**: mtDNA sampling attempt to retrieve human mtDNA from the ElMiron14 sample

| Sample | Sequenced reads | Mapped reads | Unique reads | % endo. | Q30 Reads | Dupli. rate | Cov. | C>T 5' | G>A 3' |
|---|---|---|---|---|---|---|---|---|---|
| 16333iL3 | 1556751 | 6636 | 1230 | 0.00 | 889 | 1.38 | 1.6 | 40.20 | 36.18 |
| 16333iL1 | 1815085 | 6938 | 2278 | 0.07 | 1926 | 1.18 | 6.02 | 37.71 | 37.66 |
| 16333iL4 | 1863438 | 15663 | 2517 | 0.12 | 2061 | 1.22 | 5.63 | 41.10 | 42.30 |
| 16333iL5 | 3676163 | 18377 | 2826 | 0.07 | 2237 | 1.26 | 5.4 | 38.19 | 41.41 |
| 16333iL4.1 | 3650823 | 31551 | 1344 | 0.08 | 841 | 1.60 | 0.8 | 40.71 | 42.85 |
| 16333iL4.2 | 1156483 | 11663 | 1224 | 0.12 | 781 | 1.57 | 1.16 | 35.88 | 34.96 |
| 16333iL4.3 | 2862850 | 30896 | 1514 | 0.04 | 955 | 1.59 | 1.02 | 36.41 | 32.98 |
| 16333iL4.4 | 3021893 | 12014 | 1219 | 0.05 | 771 | 1.58 | 0.90 | 35.41 | 37.16 |
| 16333iL4.5 | 1302657 | 8932 | 1161 | 0.09 | 749 | 1.55 | 0.92 | 50.36 | 36.00 |

Haplogrep predicted the Haplogroup of 16333iL1 (U4'9 (77% quality)), 16333iL4 (U4'9 (82% quality)), 16333iL5 (U2'3'4'7'8'9 (81% quality)). These results are consistent with the data coming from multiple individuals, showing a diversity of U haplogroups typical of Pleistocene populations. These results suggest multiple individuals contributed the data we analyzed (rather than a single underlying individual).

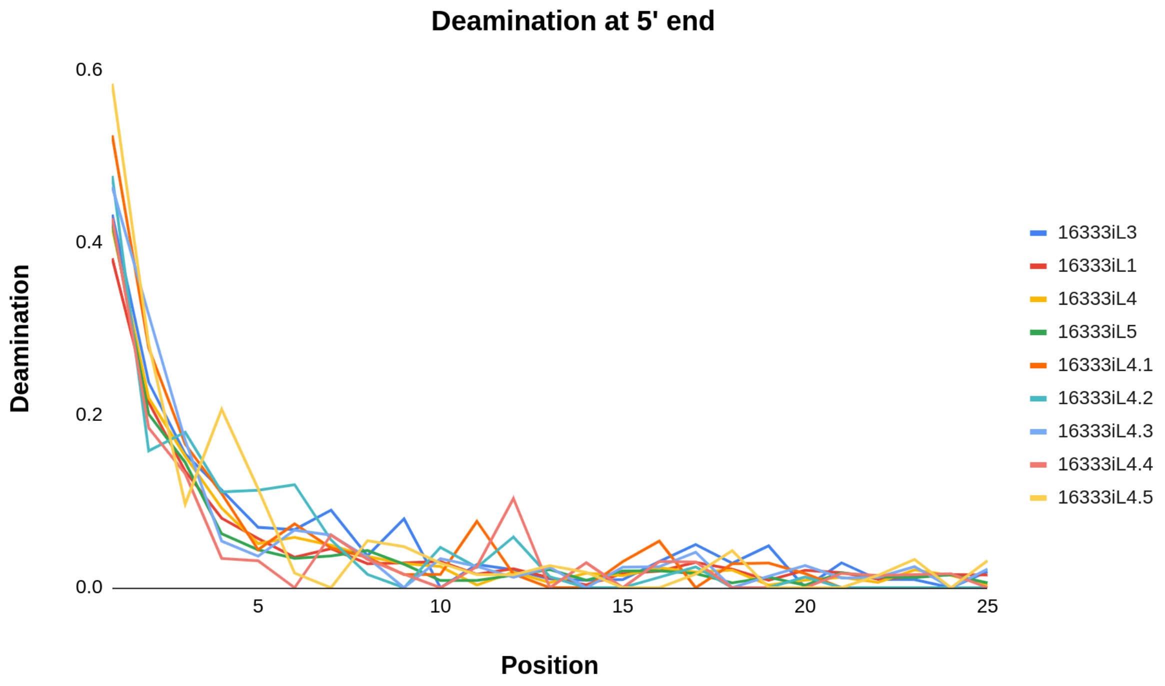


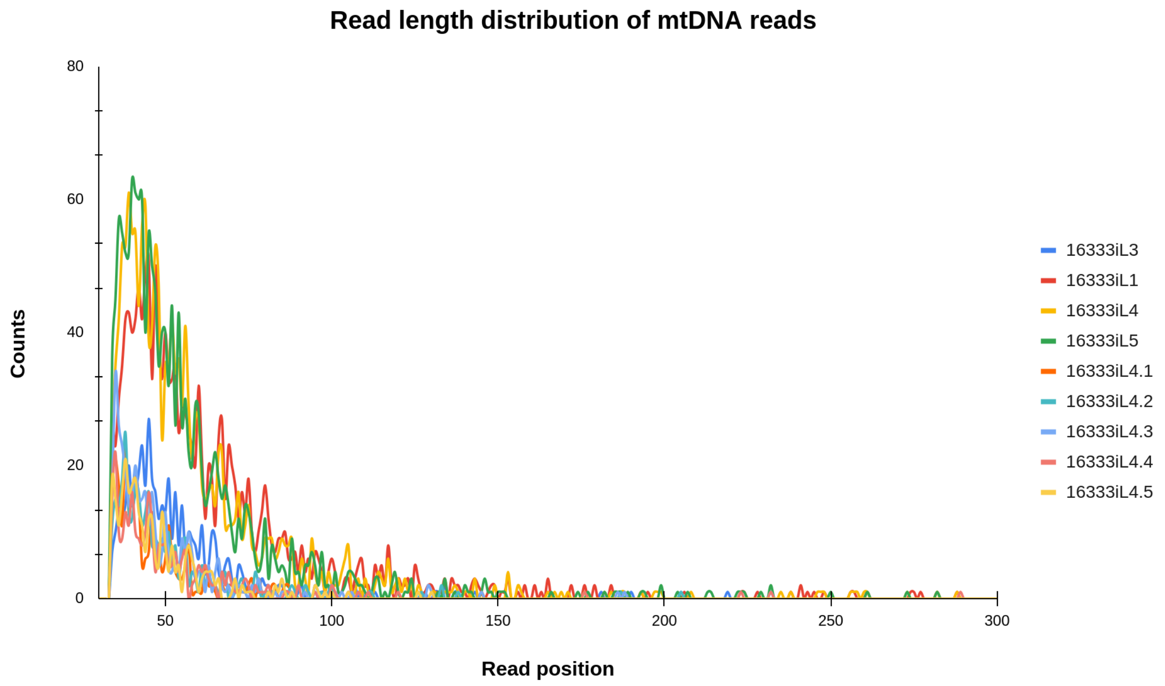


**Supplementary Figure 2:** Mitochondria DNA reads of the ElMiron14 samples, the sequenced libraries show deamination and short reads. The list of alleles is presented in Supplementary Data 2.

### 2. Chufin

#### 2.1 Sample context

The archaeological value of Chufin cave became known with the discovery of the first rock art in 1972. Immediately, the site was excavated between 1974 and 1976, and the rock art was studied [7,8]. In 2021, archaeological excavations were resumed by one of us (DGM) to define the site's stratigraphic sequence better and characterize the human occupations that took place there. Initially, two test pits were opened next to the previous excavation areas and to the rock art walls to identify intact sediments. During this first campaign, the human tooth (CHU.53k.3.02) sequenced in this study was recovered from layer 201 of the right sector of the excavation. It was located at a superficial level next to the parietal engravings and in archaeological material mainly attributed to the Solutrean period. However, the direct dating of the tooth is slightly older.

A tooth from Chufin cave (Supplementary Figure 3) was radiocarbon-dated. The sample was prepared and chemically pretreated at the Higham Lab at the University of Vienna, Austria. Collagen was extracted using a modified Longin collagen method outlined in [9] and [10]. The sample was gelatinised in weakly acidic pH3 water and ultrafiltered using a 30kD Sartorius ultrafilter, before being freeze-dried. Circa 0.3 mg of collagen was measured for $\delta^{13}C$ and $\delta^{15}N$ values using an Elemental Analyser-Isotope Ratio Mass Spectrometer (EA-IRMS) at the Faculty of Life Sciences Silver Laboratory to a precision of ±0.3‰ relative to V-PDB and AIR, respectively. Stable isotope analysis revealed a $\delta^{13}C$ value of -20.1‰ and $\delta^{15}N$ value of 11.68‰, while the atomic C/N ratio was 3.55, which suggests the collagen was of acceptable quality. The remaining collagen was combusted and graphitised with the AGE3 system. The sample was measured at the VERA (Vienna Environmental Research Accelerator) AMS facility and produced a radiocarbon age of 22793 ± 116 yr BP (VIE-1682), which is calibrated 27330-26500 cal BP, falling completely in the Gravettian in the Cantabrian region.

#### 1.1 The sequencing of Chufin tooth

We made two separate samplings from Chufin corresponding to powder 12293 and powder 15022, each of the powders was extracted once, and two double-stranded libraries were made out of each powder. We then sequenced these libraries using different methods and fractions, resulting in 46 sequencing attempts. This includes: 11 mtDNA capture samples, 13 shotgun, and 22 subsamples processed with TWIST in-solution capture. The samples captured with the TWIST capture have an absence of mitochondria reads, likely because the reagent does not target this part of the genome, as the probes were not successfully skipped (Supplementary Data 3). It is observable in the table that the reads from powder 12293 have anomalous values of typical statistics. In all the samples, the edit distance is above 2, and the deamination is around 15%. We discarded all these sequencing attempts and only processed the reads originating from powder 15022. We explored possible causes of such errors. The first analysis was conducted using the same approach as in ElMiron14 to describe the sample with BLAST and identify non-human contaminants, which were not identified. The prediction of human contamination using different software, such as Schmutzi 2.0 [11], ContamMix [12], and Calico [13], does show values around 3-5%, which could not explain this observation. From this point, we focus on the reads originating from powder 15022 (Supplementary Data 3).

#### 1.2 Chufin's Mitochondrial Genome

We recovered 261,907 unique reads aligning, after duplication removal, of the mitochondrial genome, resulting in a 1,025X genome. We explored the contamination with Calico, contamMix, and Scmutzi, first on each sequencing attempt and then on the separated libraries (Supplementary Data 3, Supplementary Tables 15-17) and then in the whole set. The merged data shows values of 0.056 (CI 0.048-0.064) with Calico and 0.02 (0.01-0.03) of contamination using Schmutzi, and finally 0.95 (CI 0.92-0.97) matching the consensus sequence (corresponding to 0.05 contamination) using contamMix.

We used the mitochondrial reads to determine the haplogroup with Haplogrep3 [14], and determined it to be U8. As the individual is female, no X contamination estimate could be performed.

Supplementary Table 15: Schmutzi contamination estimates of the studied libraries

| Library | Point est. | Down | Up |
|---|---|---|---|

| | | | |
|---|---|---|---|
| 2517 | 0.02 | 0.01 | 0.03 |
| 2518 | 0.02 | 0.01 | 0.03 |
| 2879 | 0.02 | 0.01 | 0.03 |
| 1812 | 0.03 | 0.01 | 0.05 |

Supplementary Table 16: Calico contamination estimates of the studied libraries; library 1822 did not yield results

| Library | Positions | Majority | Minority | Point est. | Lower | Upper |
|---|---|---|---|---|---|---|
| 2517 | 4 | 1240 | 59 | 0.045 | 0.034 | 0.056 |
| 2518 | 4 | 1177 | 81 | 0.064 | 0.05 | 0.078 |
| 2879 | 4 | 613 | 37 | 0.057 | 0.039 | 0.075 |
| 1812 | 0 | 0 | 0 | NA | NA | NA |

Supplementary Table 17: Contammix Contamination estimates of the studied libraries.

| Library | Point est. | Lower | Upper |
|---|---|---|---|
| 2517 | 0.03 | 0.03 | 0.04 |
| 2518 | 0.04 | 0.03 | 0.05 |
| 2879 | 0.07 | 0.05 | 0.09 |
| 1812 | 0.33 | 0.07 | 0.93 |

All the libraries showed the same mutations (against rCRS) which codify for Haplogroup U8: 73G 263G 315.1C 750G 1438G 1811G 2706G 3107C 4769G 6455T 7028T 8860G 9698C 11467G 11719A 12308G 12372A 13145A 14766T 15326G 16093C 16129A 16569A. Haplogrep predicts this haplogroup with 94% support.

We concluded that the most likely scenario is that the sample shows contamination in a range between 5% to 10%.

### 1.3 The nuclear data.

We have first analysed the sequencing libraries and identified differences between libraries and technologies. Overall, we observe significant differences in the capture-sequenced libraries' values compared to the shotgun ones. We used the differential proportion between X and Y chromosomes, 15-fold, to infer that the tooth belongs to a female.

We tested possible differences in this extract: We detect that the ED is lower in capture (mean=0.693 (SD=0.0320) vs. mean=0.765 (SD=0.0217)) Mann-Whitney U test to test the difference, *p-value*=0.002 We also detected that the deamination at the 5' end (C to T transitions) is higher in shotgun samples (mean=0.274 (SD=0.00386) vs. mean=0.158 (SD=0.00855)) with a *p-value o*f the Mann-Whitney U test of 0.001. The most significant difference is observed in the length (mean=73.5 (SD=1.15) vs. mean=57.5 (SD=2.17)), using the Mann-Whitney U test to test the difference, *p-value*=0.001. We observed a higher number of reads recovered in shotgun experiments; however, the median is higher in capture ones (Supplementary Table 18).

Supplementary Table 18: Results of the comparisons previously mentioned in the text above

| Method | Mean | Median | SD | Min | Max |
|---|---|---|---|---|---|
| Capture | 128042 | 107684 | 67117 | 52450 | 298066 |
| Shotgun | 168576 | 176984 | 25141 | 120857 | 192634 |

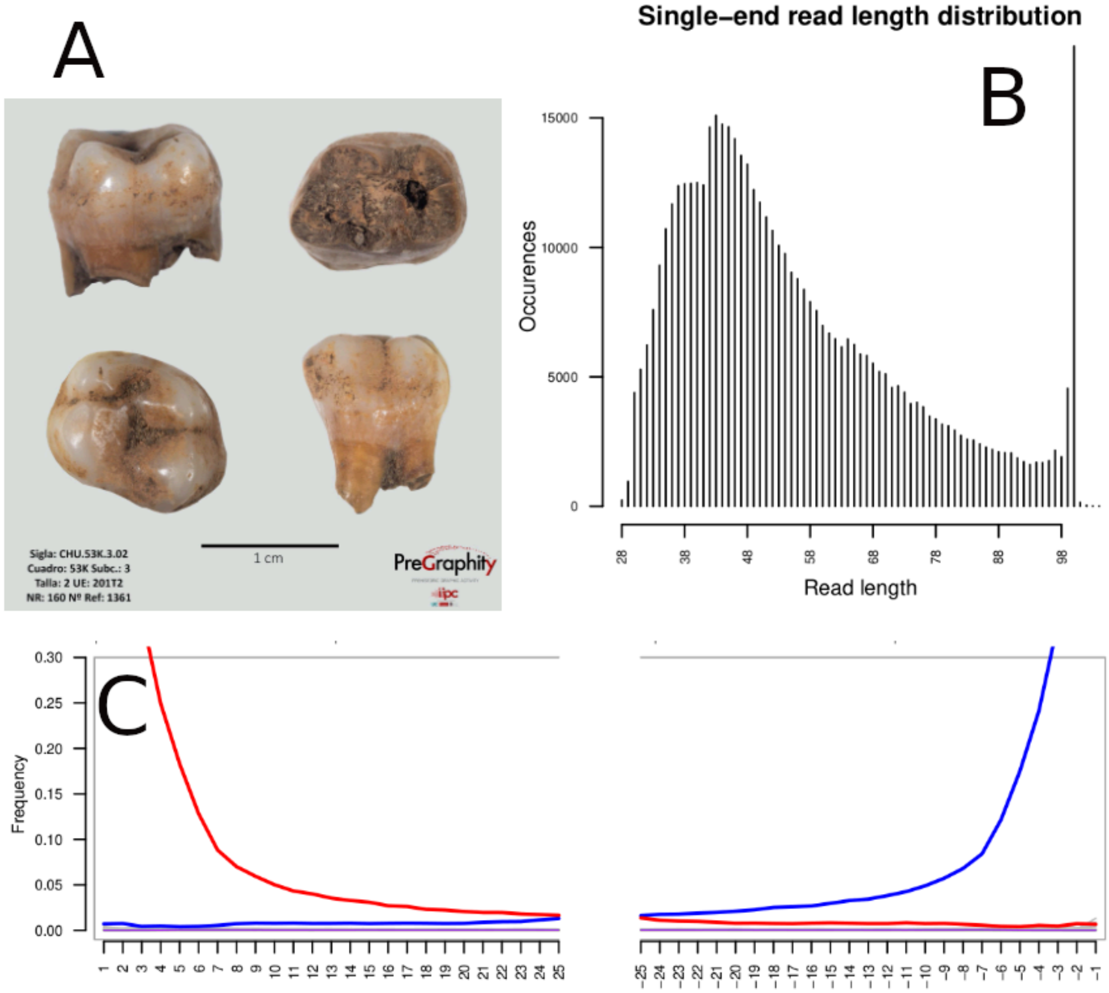


**Supplementary Figure 3:** PMD3 reads from Chufin. Bottom: Cytosine deamination patterns along sequencing reads indicate ancient DNA authenticity. Top: Distribution of read lengths, showing characteristic fragmentation patterns typical of degraded ancient DNA.

We merged the sequencing libraries from libraries 2517 and 2518, which include 27 sequencing attempts. After removing duplicates, we obtained 2,518,244 reads. We then removed duplicates between the whole set, resulting in 2,518,244, as no duplicates were found (meaning that no moleculeules were shared between libraries). We trimmed 5 bp per end following the mapDamage results using trimbam from bamUtil 1.0.15 [15]. We then called pseudo-haploid genotypes using pileupCaller from sequence tools 1.5.4.0 [4]. 141,799 SNPS were recovered in the sample with more reads.

We also approached and defined a similar strategy to the one used for elMiron14 samples, and we separately merged the Shotgun data and the Capture data, as well as the reads below 90 bp (Supplementary Table 19).

Supplementary Table 19: Results of variant calling in the four Chufin samples

| Sample | Calls |
|---|---|
| Chufin | 137237 |
| Chufin_C | 136317 |
| Chufin_S | 22026 |
| Chufin_90 | 90793 |

Based on the estimates of contamination that may point to 5 to 10% of contamination, we decided to restrict the analysis to damaged reads using PMDtools 0.60 [6]. This resulted in keeping 491,802 reads. We then clipped the last 5 bp per read end and called genotypes, which produced 26,614 SNPs of the 1240k dataset.

#### 3. Isturitz

Isturitz Cave is part of a karstic network running through Gaztelu Hill in the Arberoue Valley (Basque Country, France). Discovered in 1896, the site has been the focus of archaeological research from 1912 to the present, with a notable slowdown between the early 1960s and the mid-1990s. Human occupation at the site began during the Mousterian and extended through nearly the entire Upper Palaeolithic. The density and richness of certain archaeological layers suggest that the cave may have served as a major gathering place at various points in time, and it shows one of the most outstanding art records in SouthWest Europe[16].

In this vast cave, the Gravettian occupation has been primarily documented in the Isturitz Chamber, also known as the *Grande Salle*, covering an estimated area of around 700 square meters. Two layers were identified by early excavators. The oldest and richest, known as Ist IV and explored by R. and S. de Saint-Périer, has been attributed to the Noailles burin phase of the Gravettian. It yielded over 11,000 lithic tools and nearly 1,400 objects made from hard animal materials. However, recent studies suggest that this represents only a small fraction of the original assemblage. Nonetheless, the material clearly reflects intense and repeated human occupation over an extended period. More broadly, the Gravettian presence at Isturitz Cave appears to fall within a wide chronological range, from at least 32,000 to 28,000 calibrated years BP.

A fibula fragment IS-02A (1950-10-1) (Supplementary Figure 4) was discovered in 1950 from stratigraphic level IV. The bone is dated to 28,897-28,022 calBCE (26030±220 BP, OxAX-2616-50), a relatively late phase of the Gravettian at Isturitz. The individual is characterised to be a female of haplogroup M. We determined that this individual has a 0.019930 (SE=0.007326) Z=2.7 of Neanderthal, consistent with the rest of the individuals from the period[17]

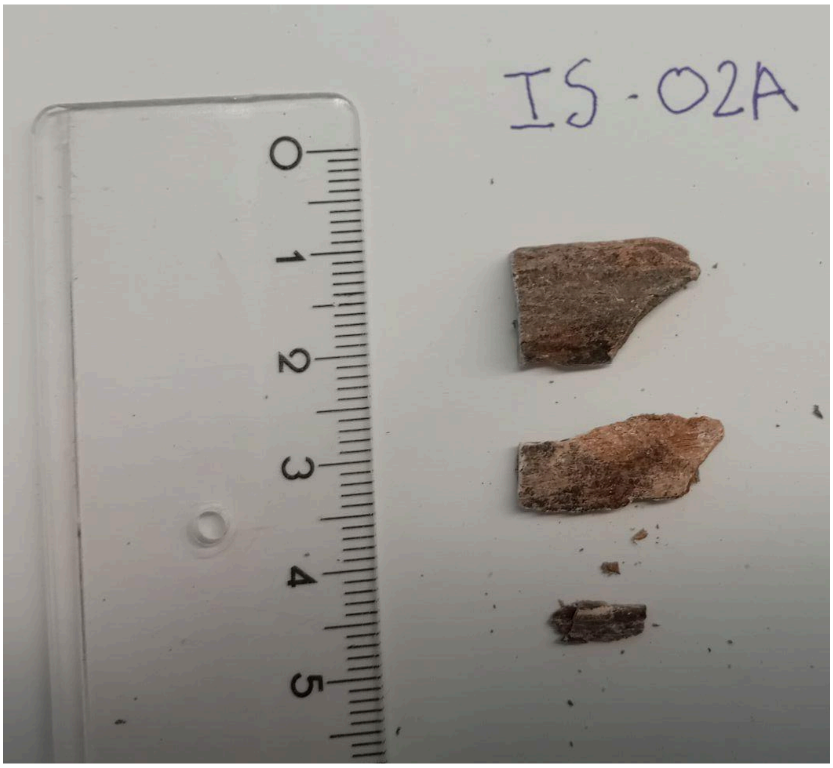


**Supplementary Figure 4:** IS-02A bone sampled in this study

4. **Ostuni1b**

The Upper Palaeolithic funerary complex of Ostuni, situated in the Santa Maria di Agnano cave (Ostuni, Apulia, Southern Italy), was discovered in 1991 by one of the authors of the present paper (DC). It yielded two primary burials, namely Ostuni 1 and Ostuni 2. Ostuni1 (Os1) individual has been dated to 27,810-27,430 cal BP and was genetically studied in Fu et al, 2016. [12]. The Ostuni 1 burial contained the exceptionally well-preserved skeleton of a young woman, who was no older than 20 years and in a late stage of pregnancy at the time of death[18,19]. The body was richly decorated with hundreds of perforated marine shells around the wrists and the cranium[18]. She was positioned on her left side in a crouched position, with the right forearm around her abdomen. Within the pelvic region of Os1, an almost complete foetal skeleton - Os1b - was recovered in excellent preservation, with the joint in connection and in the position typically acquired during the later stages of pregnancy. The Ostuni 2 burial contained a poorly preserved adult skeleton (Os2) of indeterminate sex and dated to 26339-25779 cal BC [18].

The study of the dental remains of the fetus through virtual histomorphometry has indicated that it died at 31–33 gestational weeks. The same study reports Os1b dental growth trajectories, suggesting a fetal developmental timing for Os1b slightly faster than in contemporary fetal individuals. Tree physiological stress episodes were identified in the enamel, which formed during the last two and a half months of life, suggesting that both the Os1 mother and Os1b fetus were under severe physiological stress during the last period of life. These stressors possibly resulted in the death of both the mother and the child [20].

### 5. ADMIXTOOLS 2

We used **ADMIXTOOLS 2** to identify the best-fitting admixture graph models that could explain the differences observed in the *f*-statistics for Isturitz (see Methods). Out of 106,013 generated graphs, 62 were retained as plausible models, defined as those within Δ = 2 of the best score (plausible_graphs <- combined %>% filter(score - best_score <= 2)), a commonly used threshold in likelihood-based model comparison that retains alternative models statistically indistinguishable from the best-fitting one. Within these plausible models, Isturitz was placed in a specific configuration 38 times out of 62 comparisons, while 23 models included Věstonice as a source, and the remainder represented topologies not compatible with the current demographic history.

It is important to note that Věstonice (Czechia_Gravettian) is almost the same age as Isturitz, and therefore, the signal likely reflects an unsampled ancestor rather than deriving directly from Věstonice. A binomial test confirmed that this placement was significantly more frequent than expected under a null probability of 0.5 (one-sided $p \approx 0.035$), supporting the interpretation that this topology reflects a genuine genetic signal rather than random variation. We repeated the same procedure for other individuals assigned to the Fournol cluster with more than 50,000 SNPs, Ormesson, Catalonia_Gravettian, and Malalmuerzo (Supplementary Figures 5–7).

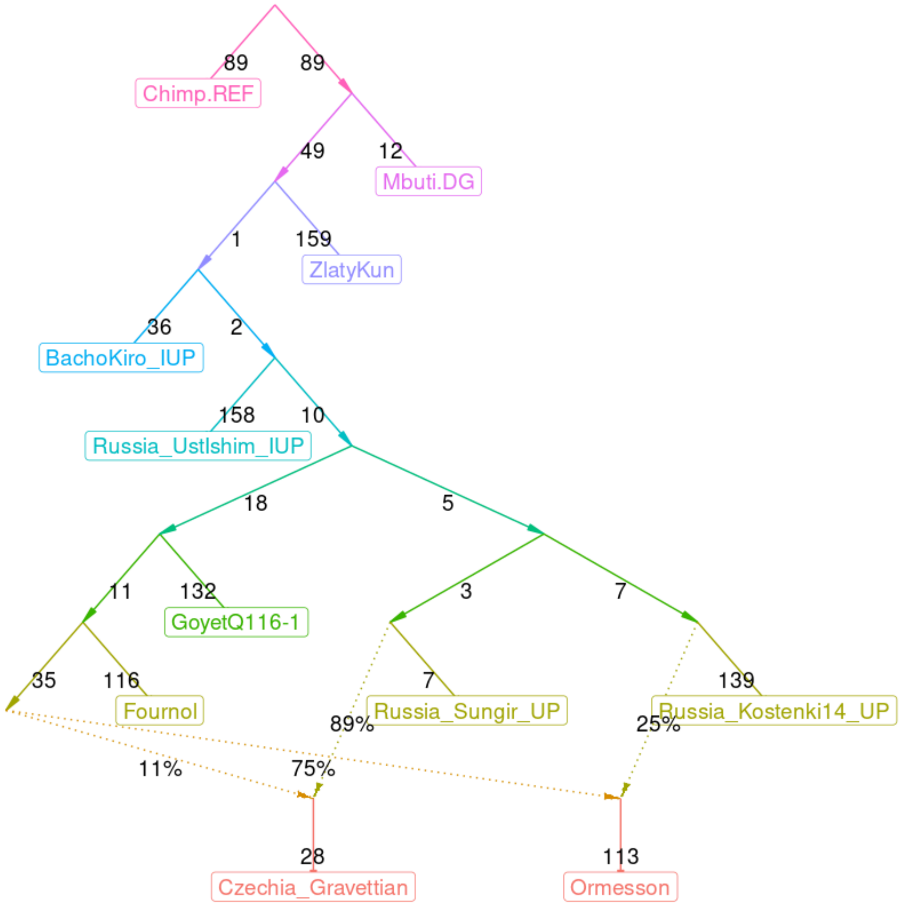


**Supplementary Figure 5:** Admixture graph fitted with the find_graphs function of ADMIXTOOLS 2, exploring models that could explain the quantitative values of the f-statistics relating the individuals. Across the 43 plausible graphs generated with these populations, Ormesson was consistently placed as an admixture of Fournol and preferably Russia_Kostirnki14_UP.

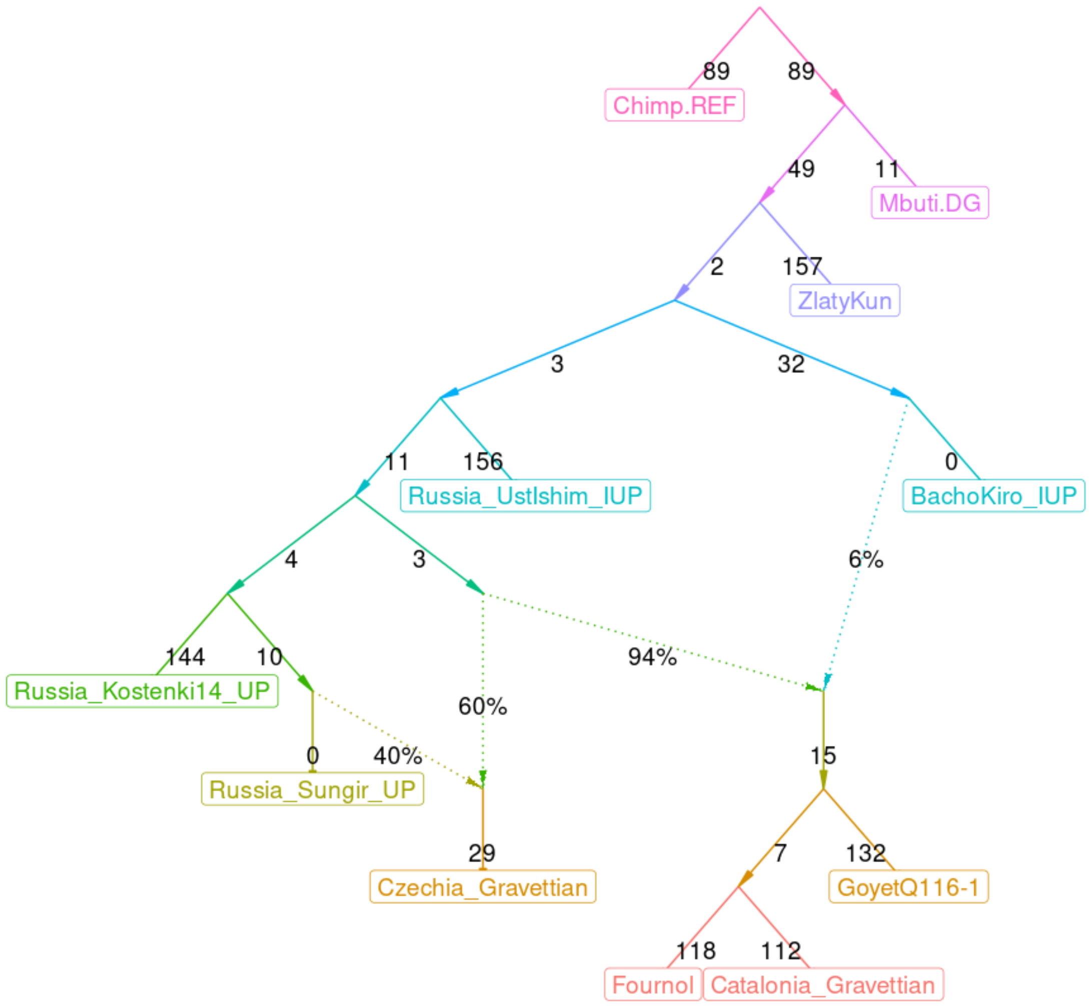


**Supplementary Figure 6:** Admixture graph fitted with the find_graphs function of ADMIXTOOLS 2, exploring models that could explain the quantitative values of the f-statistics relating the individuals. Across the 88 plausible graphs generated with these populations, Catalonia_Gravettian was consistently placed as a sister clade of Fournol.

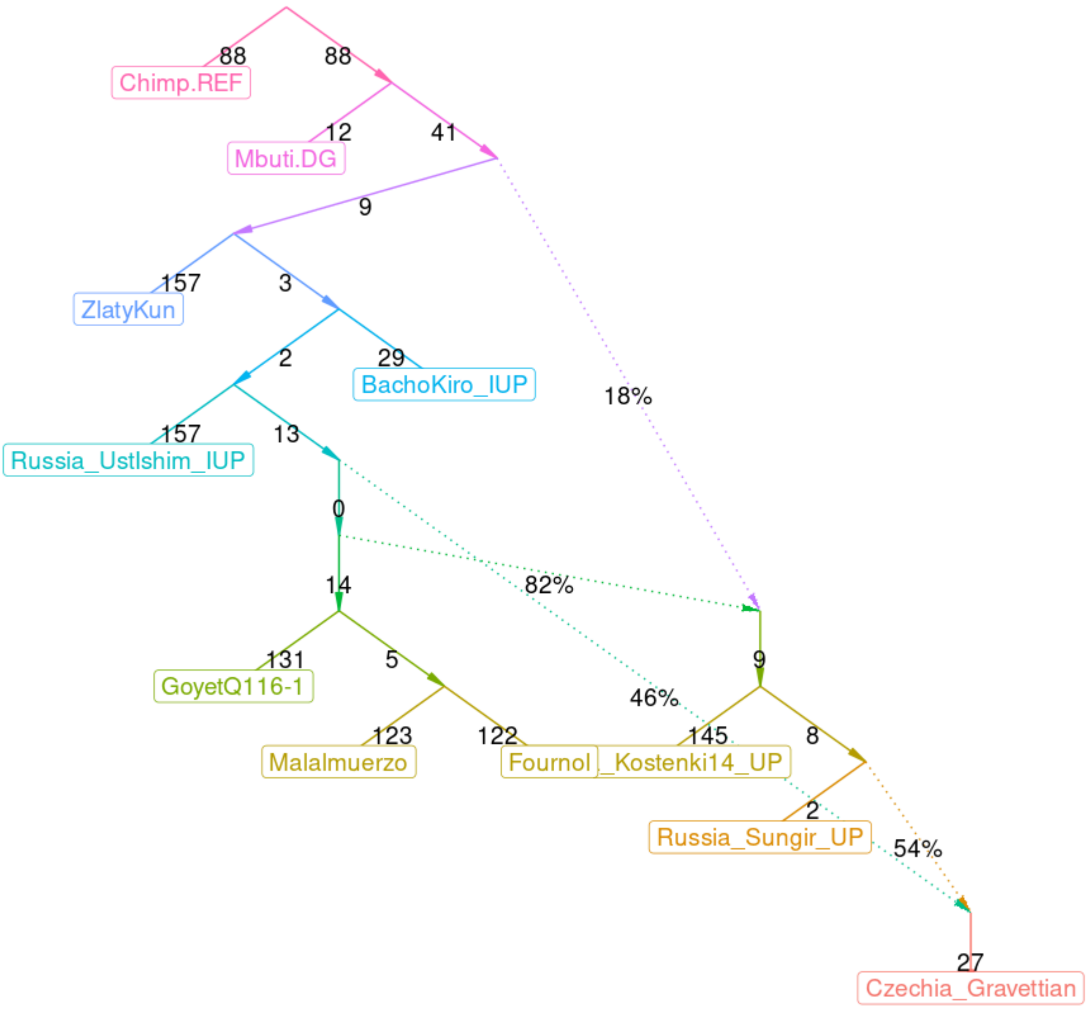


**Supplementary Figure 7:** Admixture graph fitted with the find_graphs function of ADMIXTOOLS 2, exploring models that could explain the quantitative values of the f-statistics relating the individuals. Across the 174 plausible graphs generated with these populations, Malalmuerzo was consistently placed as a sister clade of Fournol.

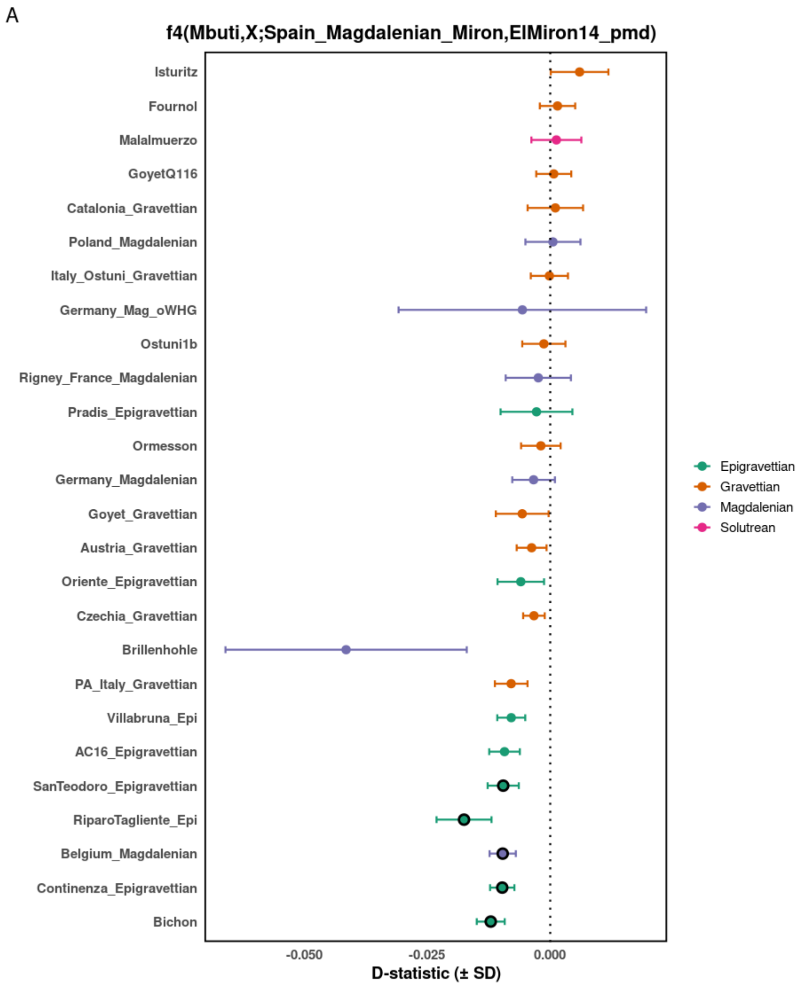


**Supplementary Figure 8:** $f_4$ comparison of individuals associated with the Gravettian, Solutrean, and Magdalenian cultures to: A) *ElMiron14* and *El Mirón_Magdalenian,* B) *Goyet Q2* and *Fournol.* Empty circles indicate non-significant statistics (−3 < Z < 3). Colours denote cultural attributions. Bars denote 1 CI.
